# Bimodal nonlinear dendrites in PV+ basket cells drive distinct memory-related oscillations

**DOI:** 10.1101/2024.09.11.612262

**Authors:** Alexandra Tzilivaki, Matthew Evan Larkum, Dietmar Schmitz

## Abstract

Neuronal oscillations are crucial for organizing distinct memory stages and behavioral states, yet the precise cellular mechanisms through which interneurons shape these dynamics remain incompletely understood. We propose and computationally explore a novel link between the nonlinear dendritic integration modes of parvalbumin-positive fast-spiking basket cells (PV+ FSBCs) and the modulation of hippocampal oscillations. Employing biophysical circuit level modeling, we test the hypothesis that PV+ FSBCs can flexibly adapt their firing patterns and influence rhythms, independent of gross changes in synaptic input quantity, by selectively engaging either their supralinear or sublinear dendritic branches. Specifically, supralinear dendrites promote high-frequency oscillations and reduce the excitation/inhibition (E/I) balance in the circuit, whereas sublinear dendrites enhance slow oscillatory power and elevate the E/I balance. This bimodal dendritic strategy provides PV+ FSBCs with an energy efficient mechanism to regulate circuit and oscillatory dynamics without necessitating large scale increases in synaptic drive. Our findings thereby uncover a previously unrecognized role for PV+ FSBC dendritic computations in regulating memory-related oscillations, offering new, experimentally testable hypotheses into the subcellular mechanisms that govern hippocampal rhythm generation.

## Main

Activity in the mammalian hippocampus is characterized by distinct frequency bands that are essential for various memory processes^1^. Neuronal activity commonly correlates with local field potential (LFP) oscillations, including slow (theta 3–10 Hz), and fast (gamma range 30– 200 Hz) bands ^1–5^. Experimental and computational work has linked these slow and fast components to different stages of memory processing and distinct behavioral states. In particular, slow oscillations are associated with memory encoding, spatial navigation, and sensory processing^4^, while fast oscillations are implicated in memory recall and consolidation ^4,6^. These components frequently co-occur through oscillatory coupling, a phenomenon considered as a signature of working memory (**Figure 1a** ) ^5,7–11^. To understand the cellular mechanisms behind these oscillatory events, significant efforts have been made to delineate the effects of various types of both excitatory and inhibitory hippocampal populations. Among them, PV+ interneurons have emerged as particularly crucial for orchestrating both slow and fast oscillations, as well as their coupling. ^9,12–17^. Nevertheless, prevailing research has largely focused on the activation or silencing of PV+ interneurons, often through optogenetic or broad computational manipulations. Consequently, the specific subcellular properties, such as dendritic processing, that can empower these interneurons to exert precise control over distinct oscillatory states and memory processes, remain largely unexplored.

**Figure 1.**
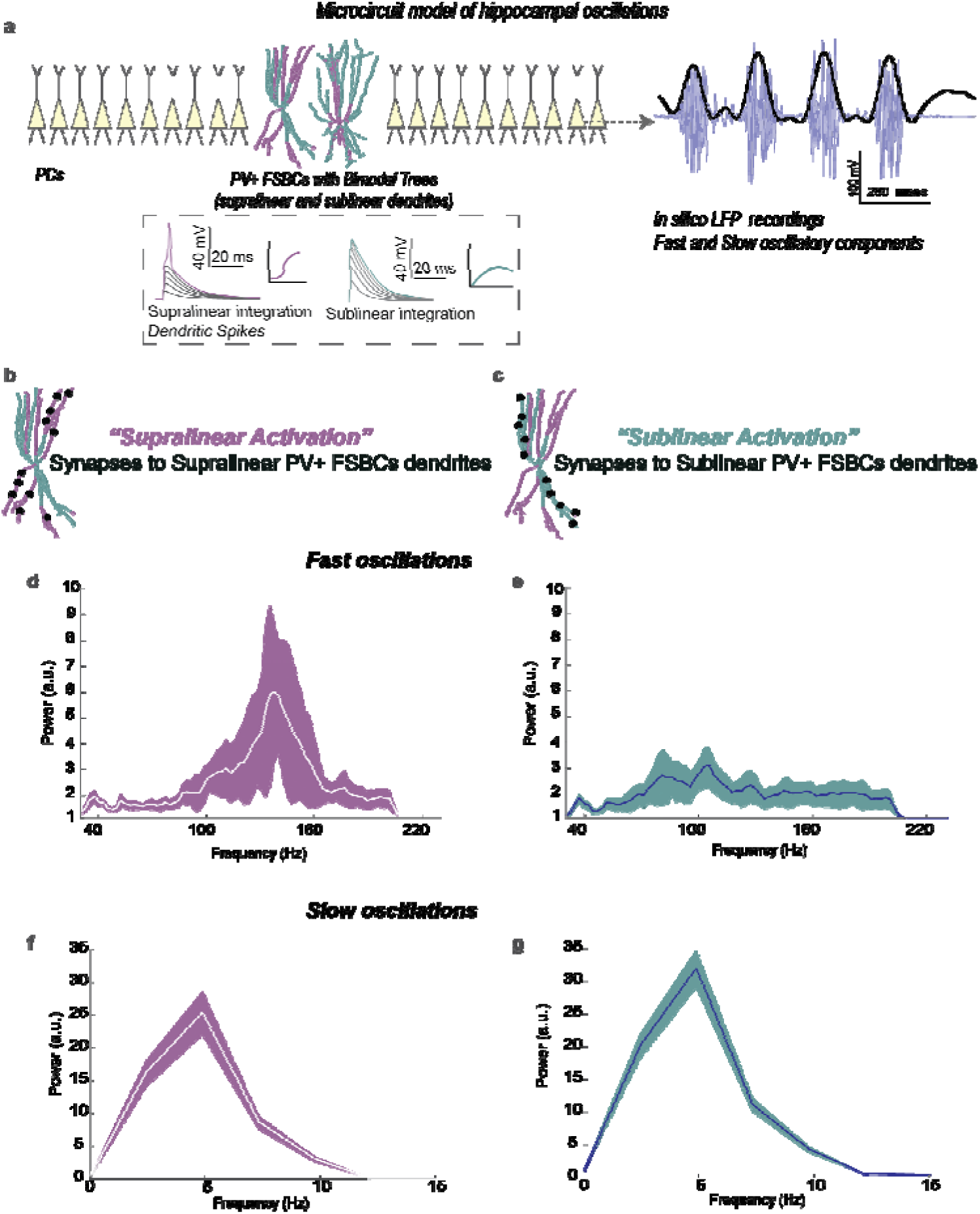
Supralinear and Sublinear branches of PV+ FSBCs trees, differentially modulate memory-related hippocampal oscillations under the same amount of synaptic activation. (**a**) Schematic Illustration of the hippocampal microcircuit model. PCs and PV+ FSBCs populations are activated by a theta (4 Hz) input for 1000 msec. An LFP electrode is positioned nearby the PCs somata. Upon activation, the model exhibits slow (bandpassed at 3-10 Hz) and fast (bandpassed at 30-200 Hz) LFP bands. PV+ FSBCs models include anatomical reconstructions and are equipped with bimodal nonlinear trees (See Glossary) as per ^33,34^. (**b-c**). The same Number of Synapses are activated either in the Supralinear (shown in purple, **b**) or in the Sublinear (shown in cyan, c) branches of the PV+ FSBCs dendritic trees. (**d-g**). Power Spectrum Density (PSD) plots under the distinct synaptic placement configurations. When only the supralinear branches receive synapses (supralinear activation), the fast oscillatory component is massively enhanced in its frequency and power compared to the sublinear activation scenario (**d,e**). In contrast, an increase in slow oscillatory power is predicted when synapses are restricted to sublinear branches (**f-g**). of the PV+ FSBCs trees. Data from 30 random simulation trials.

**Figure 2.**
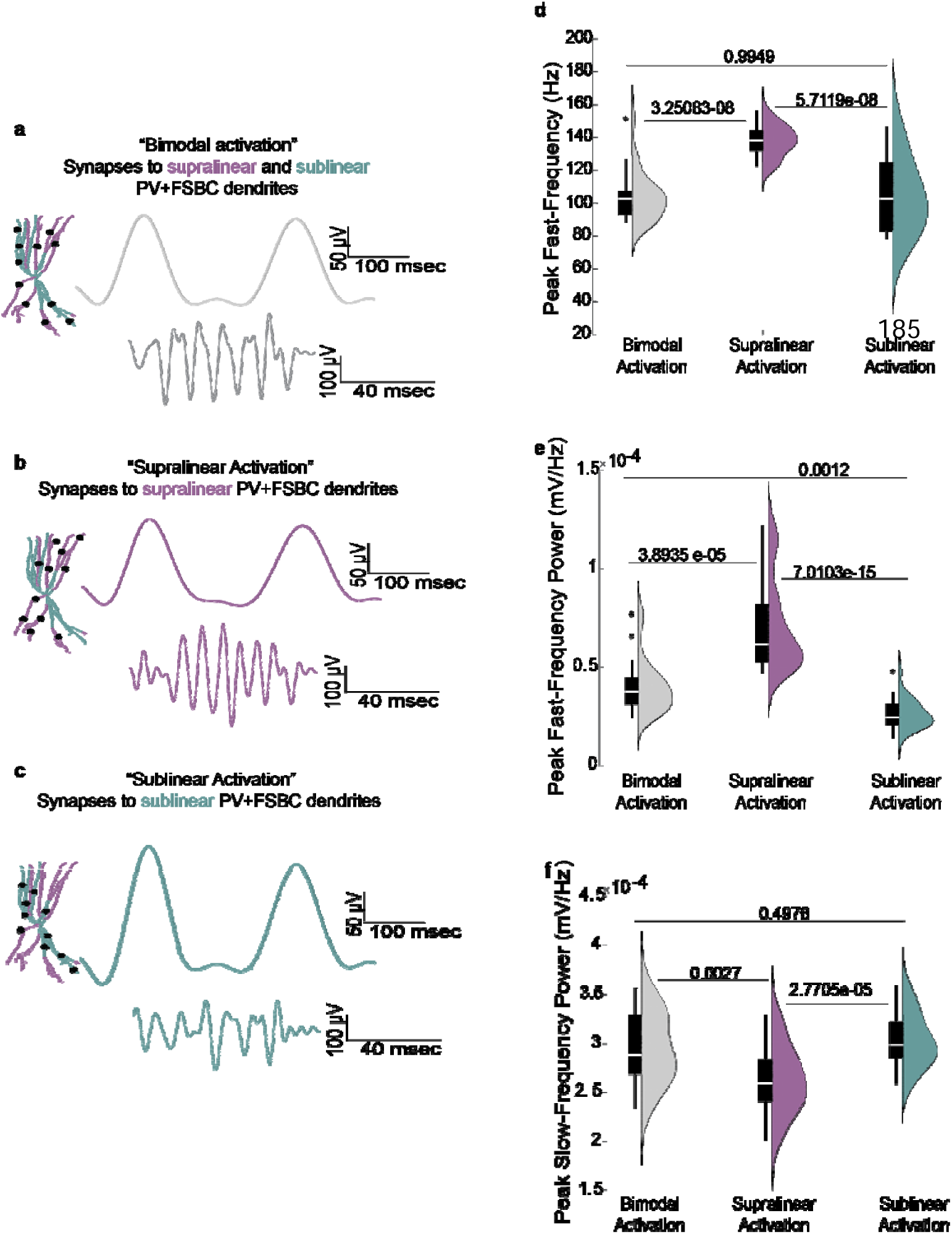
LFP frequency and power are regulated by PV+ FSBCs nonlinear dendritic activation. (**a-c**) Schematic illustration of the activation protocols and representative traces of the evoked LFP bandpassed at slow (3-10 Hz) and fast (30-200 Hz) frequences. The same number of synapses targets either both types of nonlinear dendrites (bimodal activation) (**a**), or only supralinear (**b**) or only sublinear (**c**). (**d-f**) Comparison of the peak fast-frequency (**d**) and the peak powers of the fast (**e**) and slow (**f**) LFP components. Data from 30 random simulation trials. Synaptic activation of supralinear dendrites of the PV+ FSBCs bimodal trees results in a higher peak fast-frequency and increased fast-frequency peak power, while reducing the peak power of slow oscillations compared to sublinear dendrite activation. Statistical comparisons across multiple groups were performed using the Kruskal-Wallis test followed by a post-hoc correction for multiple comparisons, suitable for data with unequal variance.

A long-standing dogma postulated that GABAergic interneurons, including PV+ cells, possess passive, purely linear dendrites incapable of nonlinear synaptic integration. This view suggested that their influence was determined only by the total quantity of synaptic inputs they receive, implying that circuit regulation during oscillations depended on aggregate synaptic drive without considering complex, potentially energy-efficient, nonlinear dendritic processing. Challenging this paradigm, recent experimental work has demonstrated that hippocampal PV+ interneurons, exhibit sophisticated nonlinear dendritic computations that significantly augment their computational power and can maximize their input-output (I/O) transformations. ^18–21^. Specifically, in an earlier *in vitro* work, Katona and colleagues reported that PV+ interneuron dendrites could express nonlinear summation^22^. Subsequent biophysical computational modeling of hippocampal and cortical PV+ FSBCs (one of the 3 major subtypes of PV+ interneurons), proposed that their dendritic trees distinctly feature coexisting supralinear and sublinear domains^23^ (**Figures 1a, S2a-d**). This predicted bimodal nonlinear processing, where PV+ FSBCs trees can leverage both nonlinear integration types, highlighted morphological and functional distinctions despite similar active properties. Supralinear branches typically possess larger volumes and lower input resistance and are capable of local generation of spikes. The smaller, high-resistance sublinear branches do not generate local spike events^23,24,26^. Notably, the core principle of this theoretical prediction, that individual PV+ neurons can possess dendrites with distinct integration rules, was later experimentally supported. Using *in vitro* 2-photon glutamate uncaging, Cornford and colleagues 21 showed the coexistence of both supralinear and (sub)linear integration within the CA1 PV+ cells whose somata locate in striatum pyramidale layer and their dendritic branches can reach striatum oriens and lacunosum moleculare layers. This experimental work, confirmed that PV+ cell dendrites are not functionally uniform, and is in strong agrrement with the modeling predictions ^23^. Thus, it further emphasized the potentially prominent role of dendritic nonlinearities in PV+ regulation in circuit dynamics across hippocampus. Although further experimental work is needed to charachterize the dendritic processing rules in other types of intereurons in the hippocampus and beyond, recent work in the L2/3 PV+ and SOM+ populations also indicates diverse nonlinear dendritic strategies^25^.

Building on these insights, the present theoretical study was motivated by fundamental questions regarding the local circuit level impact of hippocampal PV+ FSBCs dendritic nonlinearities during memory-related oscillations: How might PV+FSBCs leverage such properties to influence circuit dynamics? What advantages could bimodal, as opposed to unimodal, dendritic integration confer? And what are the distinct consequences of preferentially activating supralinear versus sublinear dendritic domains? Given the established role of PV+ FSBCs in both slow and fast oscillations, ^12,27,28^we employed advanced biophysical modeling to test our central hypothesis: that the supralinear and sublinear dendrites of PV+ FSBCs may differentially modulate their activity and, consequently, orchestrate both spiking activity and LFPs during mnemonic oscillations. Specifically, we posited that PV+ FSBCs could exploit their bimodal dendritic architecture to selectively shape fast and slow oscillatory activity within a hippocampal microcircuit (**Figures 1-4**). Such a mechanism would enable them to fine tune the E/I balance and modulate both slow and fast oscillatory signals without necessitating gross changes in synaptic input quantity (**Figures 3-4)**, thereby offering a flexible and energy-efficient strategy for engaging with distinct memory-related rhythms.

**Figure 3.**
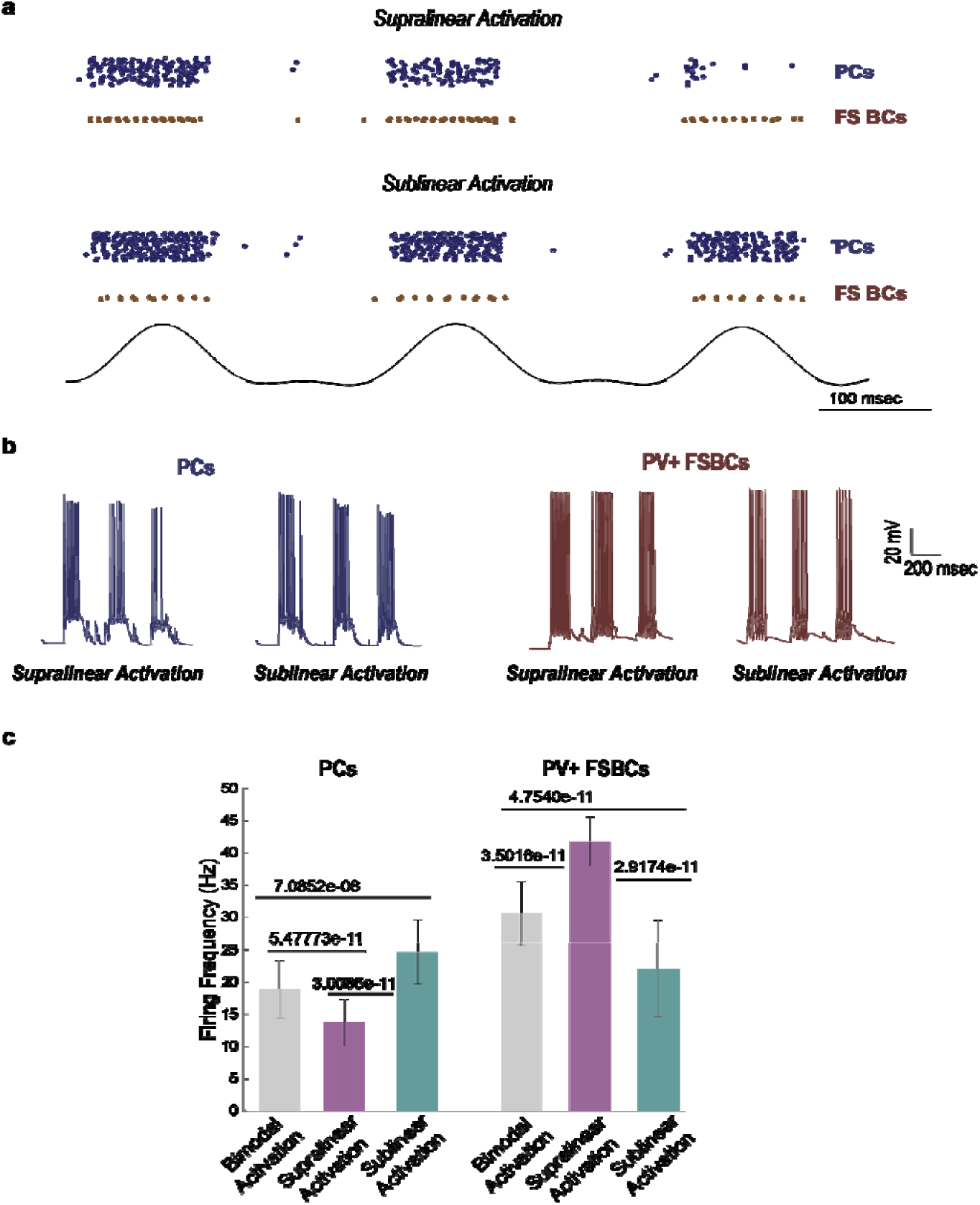
Supralinear and sublinear PV+ FSBCs dendrites differentially modulate the excitation/inhibition (E/I) balance in the microcircuit model under identical synaptic activation conditions. (**a**) Representative raster plots depicting the spiking activity of PCs (blue) and PV+ FSBCs (brown) across theta cycles when synaptic inputs are selectively targeted to either supralinear dendrites (upper panel) or sublinear dendrites (lower panel) of PV+ FSBCs. (**b**) Representative spike trains of PCs (left) and PV+ FSBCs (right). When PV+ FSBCs receive synaptic input in their supralinear branches, their firing rates increase more profoundly across theta cycles, leading to a stronger decrease in PCs spiking. Conversely, when synaptic inputs are restricted to sublinear dendrites, PV+ FSBCs firing rates are reduced, resulting in increased PCs activity. (**c**) Activation of a fixed number of synapses on supralinear dendrites (shown in purple) significantly increases the firing frequency of PV+ FSBCs, leading to a reduction in the activity of PCs when compared to bimodal (supralinear and sublinear, shown in grey) or purely sublinear activation. Conversely, activation of randomly selected sublinear dendrites (shown in cyan) decreases the firing frequency of PV+ FSBCs, resulting in higher disinhibition of the PCs. Data are based on 30 random simulation trials. Pairwise comparisons were performed using the Mann–Whitney U test.

**Figure 4.**
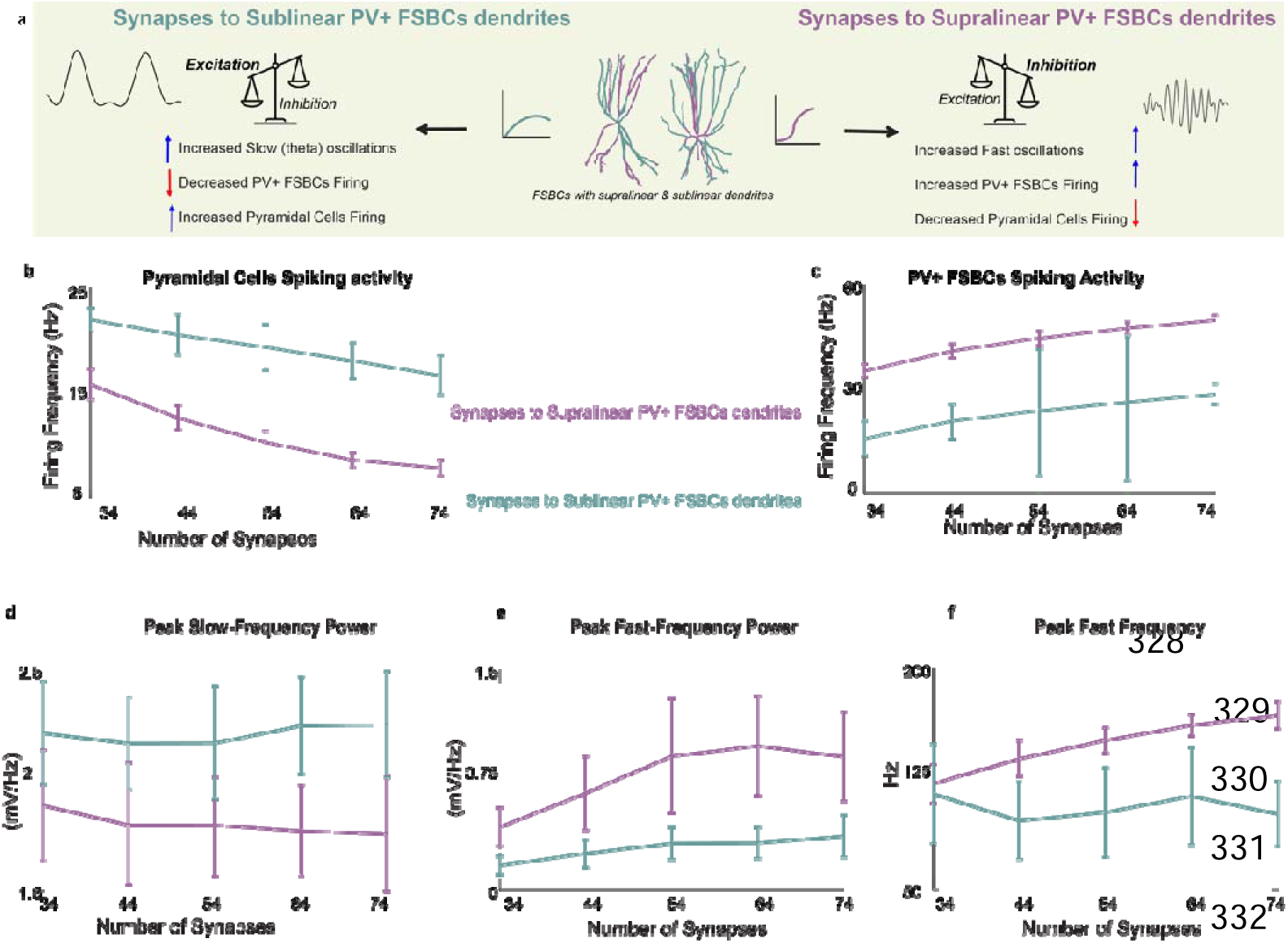
Dendritic integration mode governs PV+ FSBCs mediated modulation of hippocampal oscillations irrespectively of synaptic input amount. (**a**) Schematic summary illustrating how synaptic input targeting different dendritic integration modes in PV+ FSBCs. Sublinear (left cyan) vs. Supralinear (right purple), leads to distinct Cellular and LFP effects: Activation of sublinear dendrites reduces PV+ FSBCs firing, increases PCs activity, and enhances slow oscillations. In contrast, activation of supralinear dendrites increases PV+ FSBCs output, suppresses PCs, and strengthens fast oscillatory components. (**b–f**) Simulations upon increasing synaptic input number to either supralinear (purple) or sublinear (cyan) dendrites reveal a consistent divergence in cellular (firing frequency) and LFP responses: PV+ FSBCs firing rates increase more when their supralinear dendrites are targeted in comparison to sublinear targeting (**b**). In contrast, PCs firing is less reduced when different number of synapses are placed in sublinear PV+ FSBCs dendrites (**c**). Peak slow frequency power is enhanced with sublinear activation and reduced with supralinear input (**d**). Peak fast-frequency power and frequency is enhanced with supralinear activation and remain low under sublinear input, even when the number of synapses is high (**e–f**). Data from 30 simulation trials show mean and std values.

To test our hypothesis, we utilized an established biologically plausible hippocampal microcircuit model ^29^, that incorporates compartmental biophysical models of PCs with simplified morphologies ^29^ and PV+ FSBCs with anatomically reconstructed bimodal nonlinear dendritic trees ^23^ , mimicking the experimental phenotype of ^21^ (**Tables S1–S5, Figures. 1a, S1, S2**). Both neuronal types were extensively calibrated based on electrophysiological data (**Tables S1-S5, Figure S1**). In light of the limited experimental validation currently available for dendritic integration mechanisms in other hippocampal interneuron types (e.g., SOM+ or VIP+), and to avert speculative assumptions, here we focus in hippocampal PV+FSBCs with their characteristic bimodal nonlinear dendritic architecture (**Figure 1a**) as the sole inhibitory component within the microcircuit under investigation.

Upon activation with a theta-like input (as per ^30,31^), our model recapitulated the experimentally observed coupling of slow and fast oscillations (**Figure 1a**) ^9,10,32^, generating a slow oscillatory component alongside a fast oscillatory component (**Figure. 1a, d**). To systematically investigate how PV+ FSBCs dendritic integration modes influence network dynamics, we initially designed the following distinct activation conditions while ensuring that the total number of synaptic contacts remained constant. 1. *Supralinear Activation (shown in purple)*: To understand the effects of supralinear dendrites in these cells, we restricted all synaptic targeting exclusively to supralinear branches (**Figures. 1b,1d,1f,2b**). 2. *Sublinear Activation (shown in cyan)*: Similarly, to examine the role of sublinear dendritic processing, we constrained all synaptic inputs to sublinear branches (**Figures. 1c,1e,1g, 2c**), and 3. *Bimodal Activation* (shown in grey): Synapses onto PV+ FSBCs trees were distributed across both supralinear and sublinear dendritic branches (**Figure 2a**). This represents a mixed scenario where both types of dendrites can be targeted. In all three conditions, the total number of synaptic inputs remained the same and identical (**Figures 1-3**), and all other parameters and simulation settings were kept unchanged to ensure a controlled comparison of how supralinear vs sublinear dendrites shape PV+ FSBCs firing patterns and network oscillatory behavior. Our model predicts that supralinear dendritic activation in PV+ FSBCs preferentially amplifies the fast oscillatory component of the *in silico* LFP signals, whereas sublinear dendritic activation enhances the slow LFP component (**Figures 1-2**). Specifically, when supralinear PV+ dendrites are activated, the LFP exhibits much higher fast peak-frequency values as well as higher fast peak-frequency power values compared to sublinear activation conditions (**Figures, 1-2**). In sharp contrast, PV+ FSBCs sublinear dendritic activation significantly reduces the fast peak-frequency power relative to supralinear activation (**Figures. 1-2**), while simultaneously increasing slow LFP power values (**Figures. 1g, 2f**). These predictions suggest that PV+ FSBCs can leverage their complex dendritic architecture to selectively regulate distinct oscillatory components, dynamically shaping hippocampal network activity based on input spatial targeting rather than input amount.

To assess the robustness of the differential impact of supralinear versus sublinear dendritic activation on LFP oscillations, we performed multiple sensitivity analyses on synaptic parameters and input characteristics. In all cases, supralinear activation enhanced both peak power and peak frequency power of the fast LFP component, whereas sublinear activation consistently resulted in stronger slow oscillatory power compared to supralinear activation (**Figure, S3**).

To elucidate the mechanisms underlying the observed LFP modulations, particularly given that the total number of synapses remained constant across all three activation protocols (bimodal, supralinear, and sublinear, shown in our Figures 1-2), we analyzed the corresponding spiking activity of both PCs and PV+FSBCs (**Figure 3a-c**). Our analysis revealed that supralinear activation significantly increased PV+ FSBCs firing rates compared to both bimodal and sublinear activation scenarios (**Figure 3a, c**). This heightened inhibitory activity, in turn, suppressed PCs firing, as evidenced by the lower PCs firing frequencies observed under supralinear activation (**Figure 3b, c**). Conversely, sublinear dendritic activation diminished PV+ FSBC firing rates, leading to a disinhibition of PCs and consequently higher PCs firing frequencies relative to the supralinear condition (**Figure 3**). The bimodal activation protocol, with synapses distributed across both dendritic types, yielded intermediate firing rates for both PV+ FSBCs and PCs, indicative of a intermediate interplay between the effects of the two distinct nonlinear integration modes (**Figure 3c**). These findings collectively suggest that the nonlinear dendritic properties of PV+ FSBCs facilitate a differential modulation of network activity: supralinear dendritic inputs enhance PV+ FSBCs firing to stronger suppress PCs activity, while inputs solely to sublinear branches, result in reduced, compared to supralinear, PV+ FSBC firing, thereby disinhibiting PCs. Such dynamic regulation of the network’s E/I balance underscores the functional significance of the supralinear and sublinear nature of PV+ FSBCs trees in shaping Hippocampal circuit excitability and memory-related oscillations.

These observations demonstrate that PV+ FSBCs can dynamically adapt their firing patterns depending on whether synaptic input is received by their supralinear or sublinear dendritic branches, despite the total synaptic input remaining constant. Supralinear branches, characterized by larger volumes and lower input resistance ^33^, facilitate enhanced forward propagation of synaptic signals (**Figure S2g**). In contrast, sublinear branches, which have smaller volumes and higher input resistance ^33^, limit forward propagation (**Figure. S2f**), leading to a decrease in PV+ FSBCs firing rates. Consequently, this differential activation modulates the E/I balance in the network, directly shaping the oscillatory activity observed in the LFP. These findings indicate that PV+ FSBCs can potentially use the bimodal nonlinear trees to differentially process incoming synaptic inputs, even when the total synaptic input remains unchanged. Such a mechanism may not only modulate PV+ FSBCs firing responses at the cellular level (**Figure 3**) but also regulate oscillations **(Figure 1-2)**, thereby influencing hippocampal mnemonic functions.

It is well established that increasing the number of synaptic inputs onto a neuron can elevate its firing rate, which in turn can influence the frequency and power of oscillations. From this perspective, even sublinear dendrites, despite lacking the ability to generate dendritic spikes, can still drive high-frequency output if provided with sufficient synaptic input. Indeed, recent studies have shown that interneurons with purely sublinear dendrites can contribute to gamma oscillations when strongly activated^35^. Our model aligns with this to some extent: activation of sublinear PV+ FSBCs dendrites supports fast oscillations too (see also **Figure S4**), but their frequency and power are substantially lower compared to activation of supralinear dendrites. This raises a key conceptual question: what would be the functional advantage of PV+ FSBCs possessing both sublinear and supralinear dendrites, if either could potentially support high-frequency activity given enough input?

We propose that nonlinear dendritic diversity within the same cell may offer a computationally and energetically efficient mechanism for dynamically modulating PV+ FSBCs output and hippocampal oscillations (**Figure 4**). Activating large numbers of synapses is energetically costly. In contrast, bimodal nonlinear dendritic integration may enable PV+ FSBCs to achieve distinct functional outcomes, such as differential modulation of firing rates, E/I balance, and oscillatory states, without requiring an increase in synaptic load.

In our simulations (**Figure 4**), we systematically examine the impact of different synaptic input onto either supralinear or sublinear dendrites across multiple levels. The results show a clear dichotomy in the response depending on which dendritic type is targeted: Increasing synaptic input onto supralinear dendrites progressively enhances PV+ FSBCs firing, strongly suppresses excitatory PCs activity, and strengthens high-frequency oscillations (**Figure 4a, right side; b–f, purple curves**). In sharp contrast, increasing synaptic input onto sublinear dendrites leads to comparatively reduced PV+ FSBCs output, stronger disinhibition of PCs, and an increase in slow (theta) oscillatory power (**Figure 4a, left side; b–f, cyan curves**). Notably, even at the highest levels of synaptic input, sublinear activation fails to replicate the fast oscillatory dynamics achieved through supralinear activation. Together, these findings reveal that PV+ FSBCs can exploit their nonlinear bimodal dendritic architecture to adaptively shape cellular and circuit activity depending on input spatial location, rather than synaptic quantity alone. This supports a view in which dendritic nonlinearities may provide an energy-efficient, and flexible mechanism for interneuron-mediated control of memory related oscillatory states.

We further explored how the nonlinear integration strategies of PV+ FSBCs can impact their cellular responses and circuit effects. (**Figure. S4a-b**). Specifically, we repeated our simulations pipeline but with PV+ FSBCs models that are equipped with purely supralinear or purely sublinear trees. Leveraging the distinct morphological features of supralinear and sublinear branches in our bimodal FSBCs reconstructions, we adjusted the dendritic morphologies as per ^33^. When PV+ FSBCs were equipped with purely supralinear dendritic trees, we observed enhanced fast frequency and power compared to both bimodal and purely sublinear tree configurations. This configuration also maximized their firing (**Figure S4a, c-h**). Conversely, PV+ FSBCs with purely sublinear dendritic trees present a reduced spike rate which increased the PCs activity (**Figure. S4h**). Sublinear trees exhibited increased slow power and decreased fast power and lower fast power compared to supralinear trees (**Figure. S4b-g**). Crucially, these simulations of ‘one-mode’ integration are informative beyond direct comparison with the bimodal case (**Figures 1-4**), as they delineate the functional consequences should PV+ FSBCs, or other PV+ subtypes thereof, predominantly operate via one of these fundamental linear/nonlinear strategies, aligning with all (often diverse) experimental observations of dendritic computations in PV+ interneurons across areas^22,25^.

These simulations underscore that PV+ FSBCs may take advantage of their bimodal nonlinear dendritic trees by dynamically utilizing their spiking-supralinear or their passive linear/ sublinear dendrites to regulate the E/I balance within the network, thereby orchestrating distinct oscillatory behaviors under equivalent synaptic input conditions. Importantly, activating either only supralinear or only sublinear branches of a bimodal tree (**Figs.1-2**) exhibits similar phenotypic responses to configurations of purely supralinear or purely sublinear trees (**Fig.S3**). This further supports the notion that the bimodal nonlinear nature of PV+ FSBCs dendrites allows for flexible modulation of network activity and oscillatory patterns crucial for hippocampal mnemonic functions.

Apart from the type of dendritic integration, the spatial arrangement of synapses, along the selected dendritic branch, influences the firing characteristics^36^. It’s well established that supralinear dendrites tend to enhance clustered synaptic allocation, whereas sublinear dendrites prefer dispersed activation. For PV+ FSBCs, it has been observed that dispersed synaptic activation leads to higher firing frequencies compared to clustered ^20,33,37^. This phenomenon is influenced by the co-existence of sublinear along with supralinear branches, coupled with their small size and the presence of A-type potassium channels^33,37,38^, which discourages preference for clustered inputs. To investigate how synaptic clustering affects the oscillatory behavior of our network, we simulated the activation of the same number of synapses as in our disperse protocols (as in **Figure. S4**) but now clustered in a few randomly chosen dendrites (**Figure. S5**). As anticipated, clustered activation of sublinear PV+ FSBC trees significantly reduced their firing rates while maximizing the E/I balance in the network. Conversely, clustering synapses in supralinear trees enhanced PV+ FSBC firing rates compared to both bimodal and sublinear clustering configurations, while also decreasing the firing rates of the PCs (**Figure. S5j**). At the oscillatory level, akin to our simulations with dispersed synaptic activation, clustered synapses in supralinear PV+ FSBC trees enhanced fast LFP components, whereas synaptic clustering in sublinear trees boosted the slow LFP power (**Figure. S5a-i**). These findings underscore the dual impact of dendritic morphology and synaptic spatial arrangement on neuronal firing patterns and network oscillatory dynamics. They highlight the intricate mechanisms by which PV+ FSBCs can dynamically modulate their responses and contribute to the regulation of network excitability and oscillatory activity crucial for hippocampal function. An important consideration for future experimental and computational work is whether supralinear and sublinear dendrites receive synaptic inputs from distinct pathways which may also be temporally segregated. Such compartmentalization of inputs could profoundly influence inhibitory activity across different oscillatory and behavioral states (see Discussion).

Finally, we analyzed the Phase-Amplitude Coupling (PAC),^39^ to check the slow-fast oscillation coupling in the LFP across all cases (**Figure. S6**). Indeed, the LFP response aligns with experimental findings, indicating that slow activity entrains hippocampal networks to slow-fast coupling (**Figure S6a**)^5,8,40–44^. Activation of either supralinear or sublinear branches did not alter the coupling between the two oscillatory components (**Figure. S6b-f**). However, coupling was enhanced when synapses were clustered on fully sublinear dendritic trees of PV+ FSBCs. Conversely, coupling was reduced when similar clustering occurred on supralinear PV+ FSBCs dendritic trees (**Figure. S6g-j**). These results indicate that as the power and frequency of the fast component decrease, it becomes more synchronized with the slower component.

## Discussion

### 1. Bimodal Dendritic Integration in PV+ FSBCs: A Mechanism for Rhythm-Specific Modulation?

While the role of interneurons in memory-related hippocampal oscillations is well established ^45–48^, the subcellular mechanisms through which they influence these rhythms remain poorly understood. PV+ FSBCs, a major PV+ interneuron subtype, have recently been implicated in fast gamma and sharp wave-ripple (SWRs) oscillations^49^, and dendritic supralinearity has been proposed as a mechanism for enhancing fast activity ^50^ However, no direct mechanistic framework has previously explained how different, and possibly diverse dendritic integration modes such as the bimodal profile explored here could contribute to selective control of slow vs. fast memory-related oscillations. Our computational predictions serve in addressing a key knowledge gap by proposing a novel framework where the dendritic integration modes, rather than merely the synaptic input quantity, may constitute a critical determinant of the oscillatory regime that PV+ FSBCs support. Specifically, activating supralinear dendrites significantly promotes PV+ FSBCs firing, decreases the E/I balance, and amplifies fast oscillatory power and frequency. Conversely, engaging sublinear dendrites leads to less prominent PV+ FSBCs output, consequent disinhibition of PCs, and an enhancement of slow oscillations (as detailed across **Figures 1–4** and **Suppl. Figures 4-5**). Notably, these distinct dynamic outcomes manifest even when the total synaptic input to PV+ FSBCs is held constant (**Figures 1-3**), strongly suggesting that these interneurons possess an intrinsic, spatially defined computational mechanism for precise oscillatory tuning.

### 2. Implications for Memory Encoding, Consolidation, and Recall

Memory formation involves distinct temporal phases such as encoding, consolidation, and recall, each linked to specific oscillatory states (e.g., theta for encoding, gamma and SWRs for consolidation, theta-gamma coupling for recall). Our results suggest that PV+ FSBCs dendritic integration modes are ideally suited to switch the hippocampal network between these functional states: In one hand, sublinear dendritic activation, which enhances slow power and disinhibits PCs, may support learning and encoding, when increased excitability is beneficial for storing novel information. On the other hand, supralinear activation, which may lead to more pronounced suppression of the PCs and enhances high-frequency synchronization, may facilitate consolidation and recall, by reinforcing stored traces and refining the temporal coordination of spike timing. This dual-mode mechanism could presumably enable PV+ FSBCs to fine-tune their engagement with both rhythms depending on which type of nonlinear dendritic domain is driven. Thus, the functional value of dendritic nonlinearities in hippocampal PV+ FSBCs may not be merely electrical, but cognitive^51,52^: it could offer a subcellular lever for dynamically tuning inhibition in alignment with memory demands.

### 3. A Framework for Energy-Efficient and Dendritic Specific Computation

Beyond functional flexibility, energy efficiency emerges as a critical computational advantage. While both supralinear and sublinear dendrites can support PV+ FSBCs firing, our simulations (**Figure 4**) highlight that supralinear branches achieve fast-frequency modulation more effectively and at a lower synaptic cost. This supports the view that dendritic spikes are not essential for fast rhythms per se, (in agreement with ^35^) but may offer a more economical and spatially targeted solution. This prediction serves as a new idea with implications for how neural systems balance performance and metabolic constraints at the cellular and network level computations^51^. Importantly, these findings refine our understanding of PV+ FSBCs contributions: their nonlinear bimodal dendritic architecture can enable input-specific, rhythm-specific, and energy-aware modulation of activity at the circuit level. Such a perspective complements classical models where inhibition is adjusted by input quantity alone and opens the door to studying location-sensitive memory modulation at the dendritic level.

### 4. Future Directions: Generalizing Across Interneuron Types and Circuits

While our simulations establish a robust computational foundation for understanding PV+ FSBCs, they concurrently pave the way for broader experimental investigations. The hippocampus alone harbors over 20 distinct interneuron subtypes, each characterized by unique morphological, developmental, and connectivity profiles. Significant heterogeneity exists even within major inhibitory classes; for instance, the PV+ interneuron family, in addition to PV+ FSBCs, includes other subtypes such as the bistratified and axoaxonic cells. Each subtype exhibits substantial variations in its morphological, molecular, and intrinsic properties, as well as distinct connectivity patterns (e.g., perisomatic, dendritic, or axonal targeting) within local circuits. ^45,46^. Encouragingly, recent experimental work has begun to reveal nonlinear dendritic properties in other inhibitory populations beyond the hippocampus^25^. Nevertheless, for most hippocampal inhibitory subtypes, the specific dendritic integration modes, and the functional relevance of their input targeting remain largely uncharacterized. Our current modeling framework is designed for modularity and can be readily adapted to incorporate such emerging experimental data. As new biophysical and morphological profiles for other interneuron classes and subtypes are available, our model can be expanded. This will enable simulations to predict how diverse dendritic architectures and input targeting strategies contribute to distinct oscillatory roles across various inhibitory cell types. To facilitate and guide these future experimental endeavors, we subsequently outline several methodological approaches for validating our model’s predictions:

First, *ex vivo* whole-cell patch-clamp recordings combined with two-photon glutamate uncaging in brain slices can be employed to functionally identify distinct dendritic integration profiles in interneurons, like the approaches used in Cornford et al., 2019 for investigating the diverse (supralinear, and linear/sublinear) integration properties in hippocampal PV+ dendrites^21^. Additionally, voltage-sensitive dye imaging and two-photon fluorescence imaging could help visualize dendritic activity during synaptic stimulation, building on prior work by Katona et al., 2011^22^. *In vivo*, photoactivatable fluorescent proteins (e.g., PA-GFP or PA-mCherry), driven by interneuron-specific promoters, can be used for precise localization and targeting of dendritic subtypes. To assess dendritic function during memory-related oscillations, *in vivo* two-photon voltage imaging in awake behaving animals could reveal whether supralinear and sublinear dendrites are differentially engaged across theta and gamma rhythms. Simultaneous LFP recordings with dendritic imaging could further clarify how these dendritic domains contribute to oscillatory signatures, similar to what shown in the pioneering work of Liao et al.,^53^. Moreover, optogenetic activation of input pathways mapped to specific dendritic regions would allow causal testing of their roles in modulating interneurons firing and network dynamics. Finally, mapping input selectivity onto distinct dendrites (i.e. supralinear and sublinear) could be achieved through *in vitro* mGRASP-based methods, or *in vivo* monosynaptic rabies virus tracing. These approaches could reveal whether specific afferent sources (e.g., from entorhinal cortex, medial septum, or locus coeruleus) preferentially target distinct dendritic domains. Together with synergistic modeling efforts, such integrative strategies would offer a powerful path toward uncovering how dendritic processing in interneurons can dynamically and energy efficiently regulate the temporal architecture of memory processing in the hippocampus and potentially across broader cortical circuits.

### Limitations of the Study

Our microcircuit deliberately focuses on PV+ FSBCs, as this is the interneuron subtype with the most detailed experimental data on dendritic integration, but it does not include the full diversity of hippocampal inhibitory cells. Furthermore, our model utilizes a generic rhythmic input, as it remains an important open question whether distinct afferent pathways preferentially target the supralinear versus sublinear dendritic domains of PV+ neurons.

## Author contributions

AT and DS conceived the study. AT designed and ran the simulations, analyzed the data and prepared the figures. MEL and DS provided senior conceptual input on the study. DS funded and supervised the project. AT wrote the manuscript. All authors edited and approved the final version of the manuscript.

## Acknowledgements

A.T. was supported by the DFG with the SFB1315-2 TP A01 Brenda Milner Award and the Einstein Center for Neurosciences Berlin Fellowship. M.E.L. was supported by the European Research Council (101055340, ERC AdG, Cortical Coupling). D.S. was supported by the Einstein Foundation Berlin, the European Research Council (ERC) under the Europeans Union’s Horizon 2020 research and innovation program (BrainPlay grant, agreement no. 810580), the German Research Foundation (Deutsche Forschungsgemeinschaft [DFG], SFB-958 – project 184695641, project 431572356, FOR 3004 – project 415914819, SFB 1315 – project 327654276 and under Germany’s Excellence Strategy – Exc-2049-390688087 NeuroCure), and the Federal Ministry of Education and Research (BMBF, SmartAge – project 01GQ1420B). We thank very much Dr. Michalis Pagkalos, Dr. Nikolaus Maier, Dr. John Tukker, Dr. Daniel Parthier, Dr. Spiros Chavlis and Dr. Jeremie Sibille for critical discussions, proofreading and comments. We also thank Prof. Dr. Nelson Rebola, Dr. Yann Zerlaut, and Dr. Yiota Poirazi, for comments on an earlier version of the work.

## Code and Data availability

All codes/scripts and datasets required to reproduce the results and figures, as well as all statistical analyses, are accessible in the ModelDB database. https://modeldb.science/2018008 *Access code:* 280621Interneurons! They are also available on GitHub: https://github.com/AlexandraTzilivaki/Tzilivakietal2024 For questions contact

## Methods

### Model implementation and availability

Simulations were conducted using NEURON (v7.6)^54^ on a High-Performance Computing Cluster, utilizing 111 CPU cores on a 64-bit CentOS Linux operating system.

**All codes/scripts and datasets required to reproduce the results and figures, as well as all statistical analyses, are accessible in the ModelDB database.**

Please visit: https://modeldb.science/2018008 Access code: 280621Interneurons!

They are also freely available on GitHub: https://github.com/AlexandraTzilivaki/Tzilivakietal2024

### Neuronal populations

1. PV+ FSBCs

Two multi-compartmental biophysical models of CA3 PV+ FSBCs were employed, adopted from Tzilivaki et al. (2019)^33^. These models include detailed anatomical reconstructions of somata and dendritic trees taken from Neuromorpho database (originally published in Tukker et al. 2007^55^), (**Figures 1, S1-2**). Both FSBCs feature bimodal nonlinear dendritic branches, characterized by both supralinear and sublinear branches. They are equipped with fast voltage-dependent sodium channels (gnafin), delayed rectifier potassium channels (gkdrin), slow inactivation potassium channels (gslowin), slow calcium-dependent potassium channels (gkcain), A-type potassium channels in proximal and distal dendritic regions (gkadin, gkapin), h-currents (ghin), and L-, N-, and T-type voltage-activated calcium channels (gcal, gcan, and gcat, respectively). These models have been extensively validated against experimental data and accurately capture the intrinsic features and electrophysiological responses of hippocampal PV+ FSBCs (See original reference^33^ and Supplementary Information). The mean dendritic diameter and total dendritic length were measured for the supralinear and sublinear dendrites of the two FSBCs morphological reconstructions. Dendritic volume was calculated as per Tzilivaki et al. (2019)^33^ using the following formula:

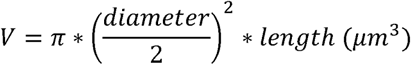

To causally manipulate the dendritic morphologies of FSBCs to generate fully supralinear or sublinear trees (**Figure S2**), we performed causal manipulations according to the approach described in Tzilivaki et al. (2019)^33^. We fixed the diameter and length of all dendrites to create trees with average supralinear or sublinear dendritic volume (**Figure S1B**), which dictated the integration mode as shown in the original modeling publication. Dendritic Input Resistance was calculated using the following formula:

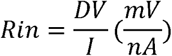

where I===−100=pA injected in each dendritic branch and DV is the generated IPSP.

For more detailed information about the models, please refer to the relevant publication^33^ and **Tables S1-2, 4-5**.

2. Pyramidal Cells (PCs)

Biophysically relevant hippocampal PC models with reduced morphology (n = 20) were adopted from Hadler et al. (2024)^29^. These models consist of somata and proximal, distal, and basal dendritic branches. PCs include a Ca2+ pump and buffering mechanism, Ca2+ activated slow AHP and medium AHP potassium (K+) currents, an HVA L-type calcium (Ca2+) current, an HVA R-type Ca2+current, an LVA T-type Ca2+ current, an h-current, a fast sodium (Na+) current, a delayed rectifier K+ current, a slowly inactivating K+ M-type current, and a fast inactivating K+ A-type current. These current mechanisms were non-uniformly distributed along the somatodendritic compartments. They were validated based on *in vitro* data to replicate the electrophysiological profile and basic dendritic architecture of hippocampal CA3 PCs^29,56^. The PC models do not include detailed nonlinear dendritic trees, as this was beyond the scope of this study. Incorporating realistic anatomical reconstructions for PCs would have significantly increased computational complexity and reduced simulation speed. For more information, see Hadler et al. (2024)^29^ and **Tables S3-5**.

### Synaptic properties

The PC models were equipped with AMPA, NMDA, and γ-aminobutyric acid type A (GABAA) synapses, while the FSBCs included Ca2+ permeable AMPA (CP-AMPA), NMDA, GABAA, and autaptic GABAA synapses. The synaptic conductance values for every connection type were calibrated based on experimental^57–60^ and modelling studies^33^ and are listed in **Table S5**

To ensure the robustness of our finding, we repeated the simulations upon increased or decreased synaptic conductance values (15% change) (see sensitivity analysis **Figure S2**)

### Hippocampal Microcircuit

To study the effect of PV+ FSBCs dendritic processing in circuit level function, we implemented an experimentally constrained microcircuit model of the CA3. Typically, for such research questions, small microcircuit modelling is the best choice as it provides enough detail to study subcellular mechanisms at the circuit level while keeping the computational demands low. Our hippocampal microcircuit configuration was adopted from Hadler et al. (2024)^29^. The model consists of 20 PCs and 2 FSBCs. In each random simulation trial (n = 30), each PC contacted up to seven (7) randomly chosen PCs with one AMPA and one NMDA synapse activation per contact. Each PV+ FSBC received synaptic input from fifteen (15) randomly chosen PCs in each simulation trial. Additionally, each PC received thirteen (13) feedback inhibitory GABAergic inputs from each PV+ FSBC per simulation trial. Each FSBC formed five (5) GABAergic synapses per simulation trial and was self-inhibited through autapses. For further details, see the Simulation Paradigms chapter and Hadler et al. (2024)^29^.

### Simulation Paradigms

Input

The microcircuit is activated by a theta entrained presynaptic population as per Turi et al., 2018^61^. The input was modeled as an artificial presynaptic population (N=22) using NEURON’s VecStim function. The spike times of the presynaptic population were generated using a sinusoidal theta like filter that was applied so to account for theta like modulated spike times.

Thata like probability formula (as per ^61^):

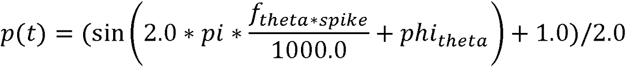

Where:

- spike = the spike time in msec (float)
- f_theta = theta-cycle frequency in Hz. (float)

For the simulations shown in **Figures 1-4** and **Supplementary** Figures 4-6, f_theta=4 Hz. For the sensitivity analysis simulations as shown in **Supplementary** Figure 3, we changed the input the theta frequency to f_theta=5Hz.

- phi_theta= theta cycle phase in radians.

For the simulations shown in **Figures 1-4** and **Supplementary** Figures 4-6, phi_theta=0 (equal to 0 radians). For the sensitivity analysis simulations as shown in **Supplementary** Figure 3, we shifted the input phase so phi_theta=0.5 (equal to 180 radians).

If the probability p(t) is greater than 0.7 a spike is generated for the specific artificial neuron (n=22). Every input neuron has its own theta modulated spike train.

Each PC received input from 5 artificial presynaptic neurons, while each PV+ FSBC received input from 7 artificial presynaptic neurons. Although the input was sufficient to activate the PC population due to its reduced morphology, 7 artificial presynaptic neurons were insufficient to evoke spiking activity in the PV+ FSBC models, given their realistic complex anatomical reconstructions. This subthreshold activation was selected to ensure that PV+ FSBCs were primarily engaged in the network due to the local inputs they received from PCs.

**1. Simulations**

The microcircuit model was simulated for approximately 12,000 milliseconds (ms) with a time step of 0.1 ms. The first 200 ms were excluded from the analysis to allow the model to reach an equilibrium state. For the dispersed protocols, synapses to FSBCs (both input and local PC-to-FSBC synapses) were randomly assigned to dendrites, meaning one randomly chosen dendrite received one pair of NMDA and CP-AMPA synapses. For the clustered protocol (**Supplementary Figure S4**), the same total amount of synapses was placed in 4 randomly chosen dendrites (cluster size: 5-7 synapses (per dendrite) Data represent the results (mean and standard deviation values (std)) of thirty (30) random simulation trials for each protocol. In every trial, the total number of synaptic contacts and the connectivity ratios remained identical, but different randomly chosen neurons (from both PC and PV+ FSBC populations) were connected to different random neurons. Additionally, in each trial, different dendrites from both PCs and PV+ FSBCs were randomly chosen and synapses were activated at different parts of the chosen dendrites. This approach ensured that the results reflected the diversity, especially of the PV+ FSBCs trees, in bimodal activation protocols. Furthermore, two different anatomical reconstructions were used for our models to account for PV+ FSBCs morphological variability (see relevant chapter).

**2. LFP simulation and Spectral analysis**

To record the Local Field Potential (LFP), an *in silico* electrode was simulated based on NEURON’s extracellular function, following the modelling approach of Vladimirov et al. (2013)^62^. The electrode was placed close to the PCs somata and remained in the exact same position throughout the execution of all protocols and simulation trials. The sampling frequency was set at 10 kHz. Despite the microcircuit model’s small size, it efficiently generated fast and slow oscillations comparable to experimental observations^5,10,39,42^ while maintaining low computational complexity and demands.

The *in silico* LFP datasets were band-passed at two respective bands: slow (3 – 10 Hz) and fast (30-200 Hz). Slow and fast Peak-Frequency powers and peak Fast frequency were determined using custom-made MATLAB scripts (MATLAB, The MathWorks Inc., Natick, MA; Torrence and Compo 1998) utilizing the p-welch function (0.4 Hz resolution) and visualized as Power-Frequency plots. Phase-Amplitude Coupling (PAC) analysis was conducted using Comodulogram generation and the calculation of the modulation index (MI) metric that were adopted from Tort et al. (2008)^32^. Wavelet phase was calculated at 15 levels from 1-15 Hz, and the amplitude at 50 levels from 30-180 Hz. The MI was obtained by measuring the divergence of the observed amplitude distribution from the uniform distribution.

### Statistical Analysis

Statistical analyses for multigroup comparisons were performed using the Kruskal-Wallis test, followed by a post-hoc correction for multiple comparisons (multcompare function Matlab). For pairwise comparisons where data exhibited unequal variance, p-values were calculated using the Mann-Whitney U test.

## Supplementary Information

**Figure S1.**
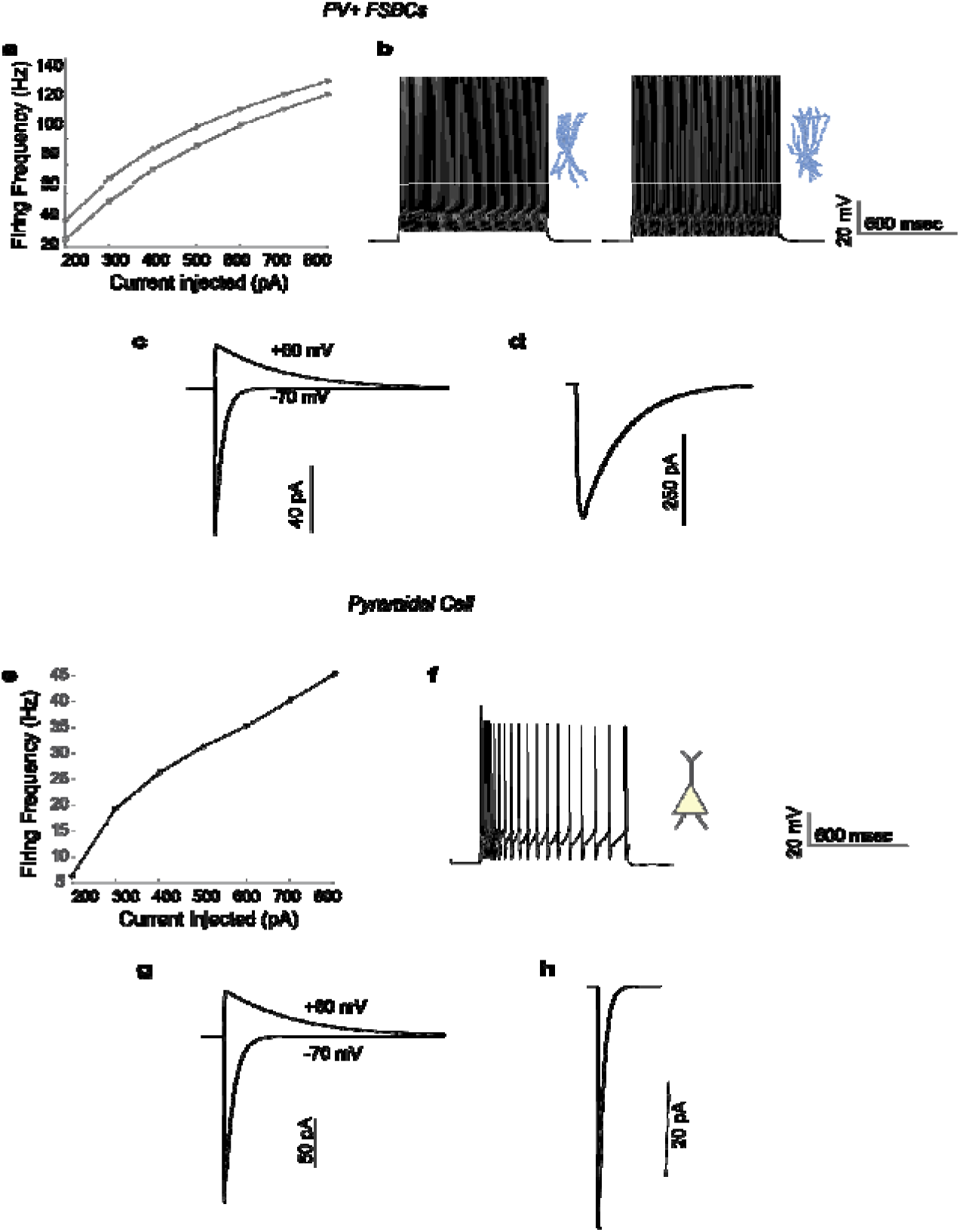
Electrophysiological calibration and responses of the models. a-c. FS BCs responses. **a.** Frequency-Injected current (FI) curve upon current clamp simulation in the cell bodies of the two multicompartmental FSBCs (1 sec duration). **b**. Firing profiles of the two FSBCs after depolarizing current injections in the cell bodies (300 pA, 1 sec). FSBCs exhibit the typical high-frequency firing pattern. **c.** CP-AMPA (−70 mV) and NMDA (+60 mV) currents upon stimulation as per ^60^. Traces represents the mean of the two multicompartmental FSBCs. **d.** A three-step voltage clamp of voltage changes from −70 mV to 10 mV (duration 1 msec) and back to −70 mV was used to produce inhibitory current. During the validation of this current, the reversal potential of Cl− was adjusted from −80 to −16 mV, in order to reproduce the experimental set up of ^63^, Mean trace of the two FSBCs responses. **e-h** Pyramidal Cell responses. **e.** FI curve of the Pyramidal Cell model same approach as in a. **f.** Firing profile of the Pyramidal Cell under current injection (300 pA for 1 sec) at the cell body. The model successfully represents the typical experimental phenotype shown in ^56^. **g.** AMPA (−70 mV voltage clamp) and NMDA (60 mV voltage clamp) currents of the Pyramidal Cell model mimic the experimental data of ^58^. **h.** Inhibitory current calibration based on experimental data from ^64^.

**Figure S2.**
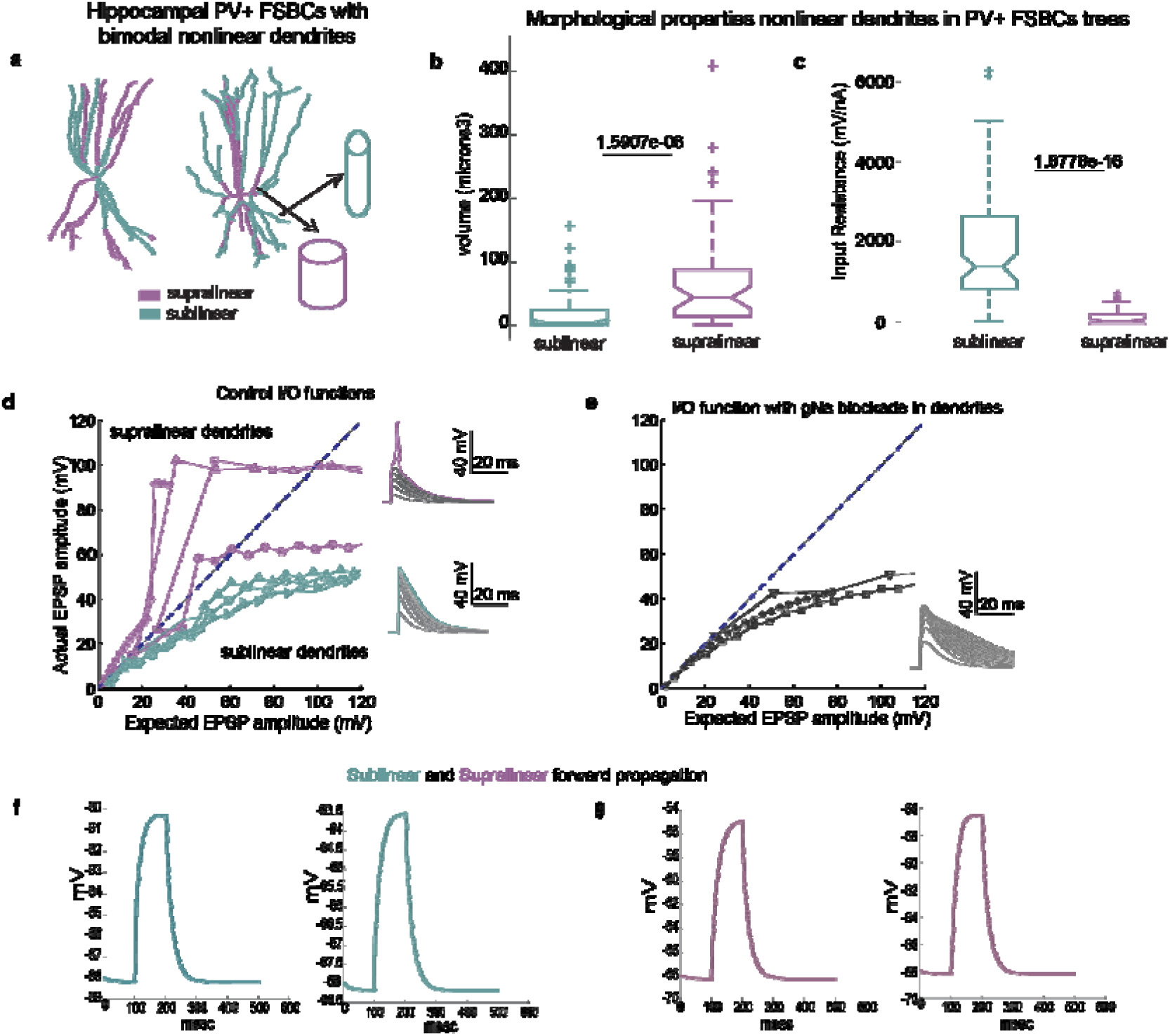
Mechanisms of bimodal nonlinear dendritic integration in Multicompartmental Models of Hippocampal FSBCs. **a**.Illustration of the morphological characteristics of supralinear and sublinear dendrites in bimodal FSBC models. Supralinear dendrites (purple) are larger, whereas sublinear dendrites (blue) are longer and thinner. **b-c**. Discriminative features between dendrite types: supralinear dendrites have larger volumes (b) and lower input resistance (c) compared to sublinear dendrites. Statistical significance was determined using the Mann–Whitney U test. **d-e**. Supralinear dendrites (purple) can generate local sodium-dependent spikes, whereas sublinear dendrites (blue) cannot (d). Blocking active sodium conductance in FSBC dendrites completely abolishes the supralinear mode (e). **f-g**. Forward propagation efficiency in bimodal nonlinear dendrites of FSBCs: Current injection (100 pA) at randomly selected dendrites and recording at the soma show that sublinear branches (f) propagate signals less effectively compared to supralinear branches (g). Panels a,d,e were adopted from ^33^

**Figure S3.**
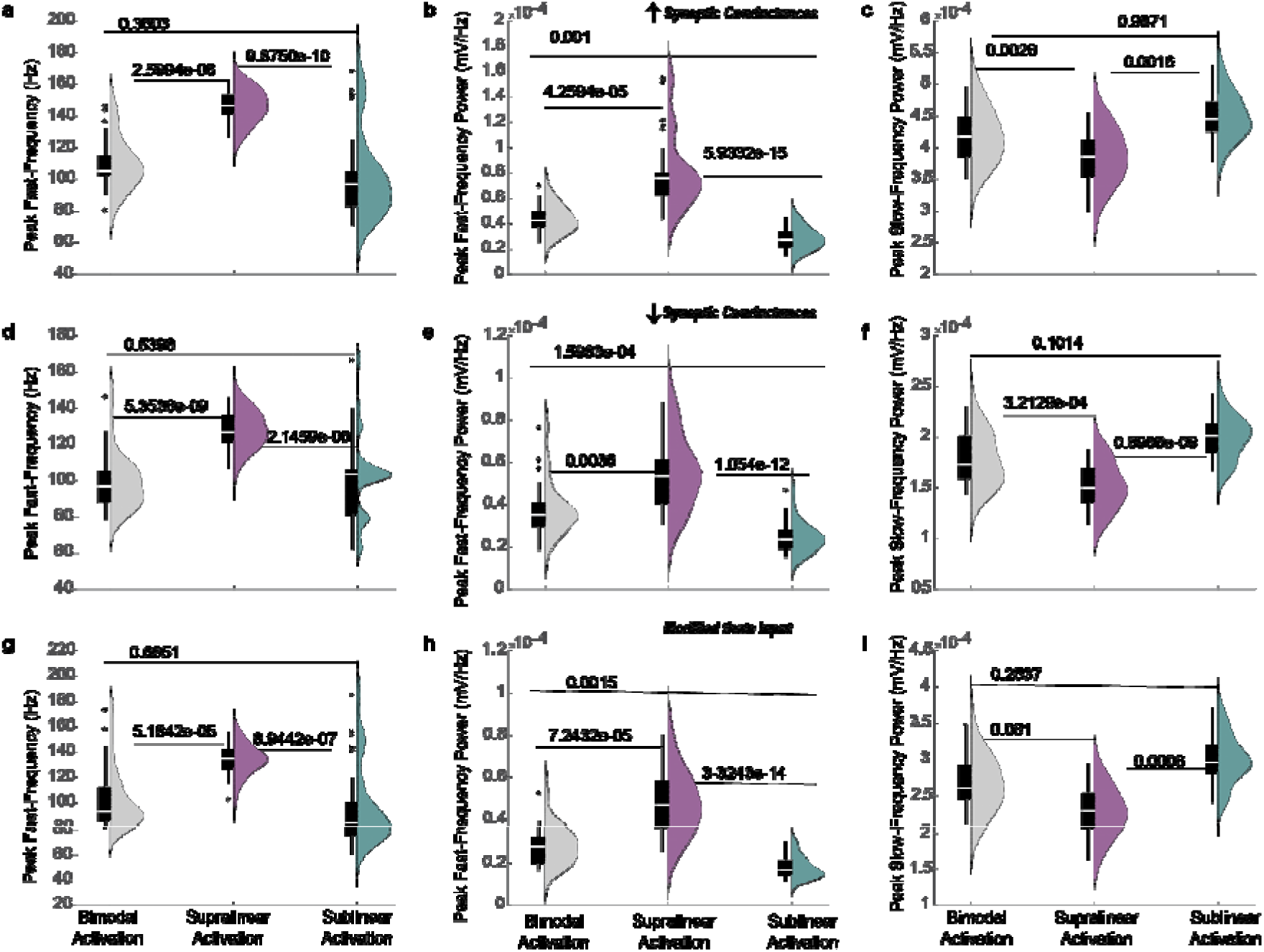
**Robustness/Sensitivity Analysis on Synaptic and Input Parameters. a-c**. 15% increase in the synaptic conductance values (applied to both input and network synapses for/from both PCs and FSBCs) do not alter the enhancement of fast peak frequency and power observed with supralinear activation. **d-f**. 15% reduction of the synaptic conductance values (for both input and network synapses involving PCs and FSBCs) also maintain the observed increase in fast peak frequency and power upon supralinear activation. **g-i**. Modifications to input parameters (input phase shifted 180°, peak frequency 5 Hz) similarly do not affect the enhancement of fast peak frequency and power driven by supralinear activation. Statistical analyses for multigroup comparisons were performed using the Kruskal-Wallis test, followed by a post-hoc correction for multiple comparisons.

**Figure S4.**
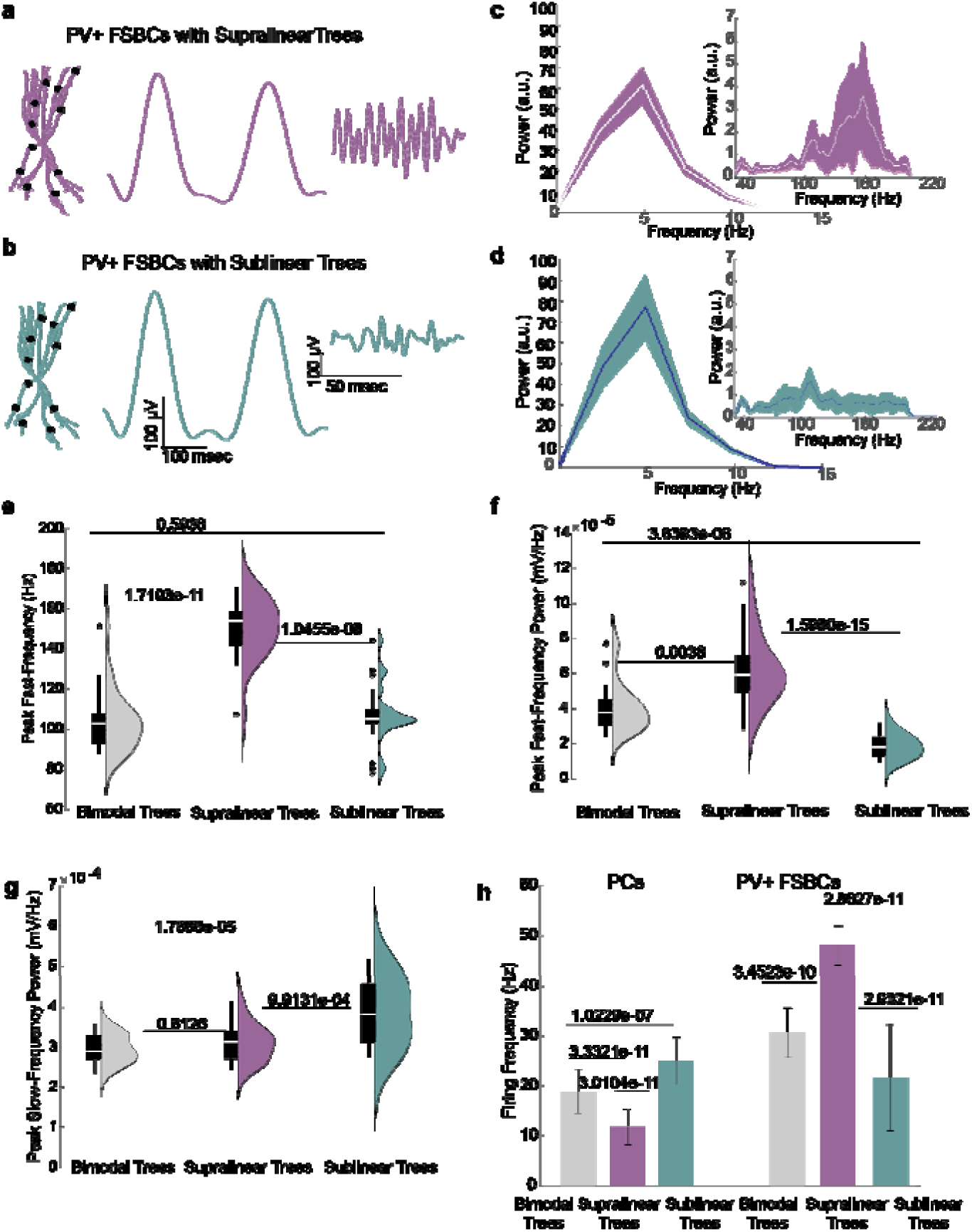
Differential Modulation of Slow and Fast LFP Components by Supralinear and Sublinear FSBCs Dendritic Trees. **a**. Activation of FSBCs equipped with purely supralinear dendritic trees, showcasing representative LFP traces bandpassed at slow (3-10 Hz) and high (30-200 Hz) frequencies. **b**. Similar to **a** but displaying activation of FSBCs with purely sublinear dendritic trees. **c-d**. Power Spectrum Density (PSD) plots of the LFP evoked when FSBCs are equipped with purely supralinear (c) or purely sublinear (d) dendritic trees, highlighting differences in frequency response. **e-g**. Comparative analysis of the peak fast-frequency (e) and peak power of fast (f) and slow (g) oscillations for FSBCs with bimodal (grey), supralinear (purple), or sublinear (blue) dendritic trees. Data are derived from 30 random simulation trials. **h**. Firing activity of the PCs and FSBCs populations within the microcircuit network across 30 random simulation trials. Activation of supralinear FSBC dendritic trees results in a decreased E/I balance compared to both bimodal (control) and sublinear trees. Statistical analyses for multigroup comparisons were conducted using the Kruskal-Wallis test followed by a post-hoc correction for multiple comparisons. Paired comparisons and p-values were calculated using the Mann-Whitney U test for data with unequal variance.

**Figure S5.**
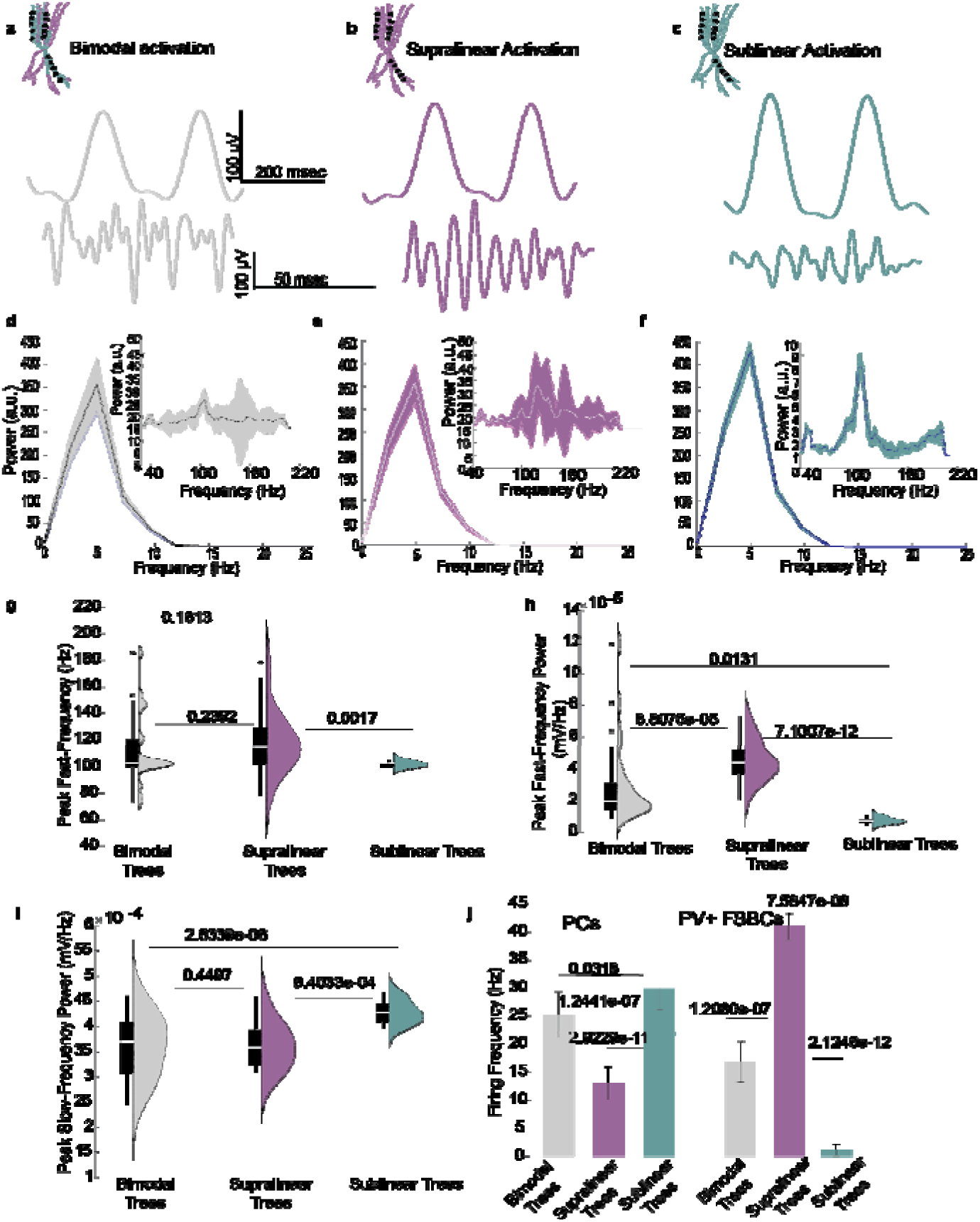
Impact of Clustered Synaptic Activation on LFP and E/I Balance in Bimodal, Supralinear, and Sublinear FSBC Dendritic Configurations. **a.** Activation of FSBCs with bimodal nonlinear dendrites (control configuration), showing representative LFP traces for slow (3-10 Hz) and fast (30-200 Hz) frequencies. Synapses are clustered in a few randomly chosen supralinear or sublinear branches. **b**. Activation of FSBCs with purely supralinear dendritic trees, showing LFP traces under clustered synaptic activation. **c**. Activation of FSBCs with purely sublinear dendritic trees, similar to conditions in a and b, illustrating the effect of clustered synaptic activation on LFP traces. **d-f**. Power Spectrum Density (PSD) plots for LFPs under clustered synaptic conditions in FSBCs with bimodal (d), supralinear (e), and sublinear (f) dendritic configurations. **g-i**. Comparison of peak frequency (g) and peak power for the fast LFP component (30-200 Hz) (h), and peak power for the slow LFP component (i), across dendritic configurations. **j**. Firing activity of PC and FSBC populations in the microcircuit network from 30 random simulation trials. Clustering in supralinear FSBC dendritic trees decreases the E/I balance in the network compared to bimodal (control) or sublinear trees. Statistical analyses for multigroup comparisons were conducted using the Kruskal-Wallis test followed by a post-hoc correction for multiple comparisons. Paired comparisons and p-values were calculated using the Mann-Whitney U test for data with unequal variance.

**Figure S6.**
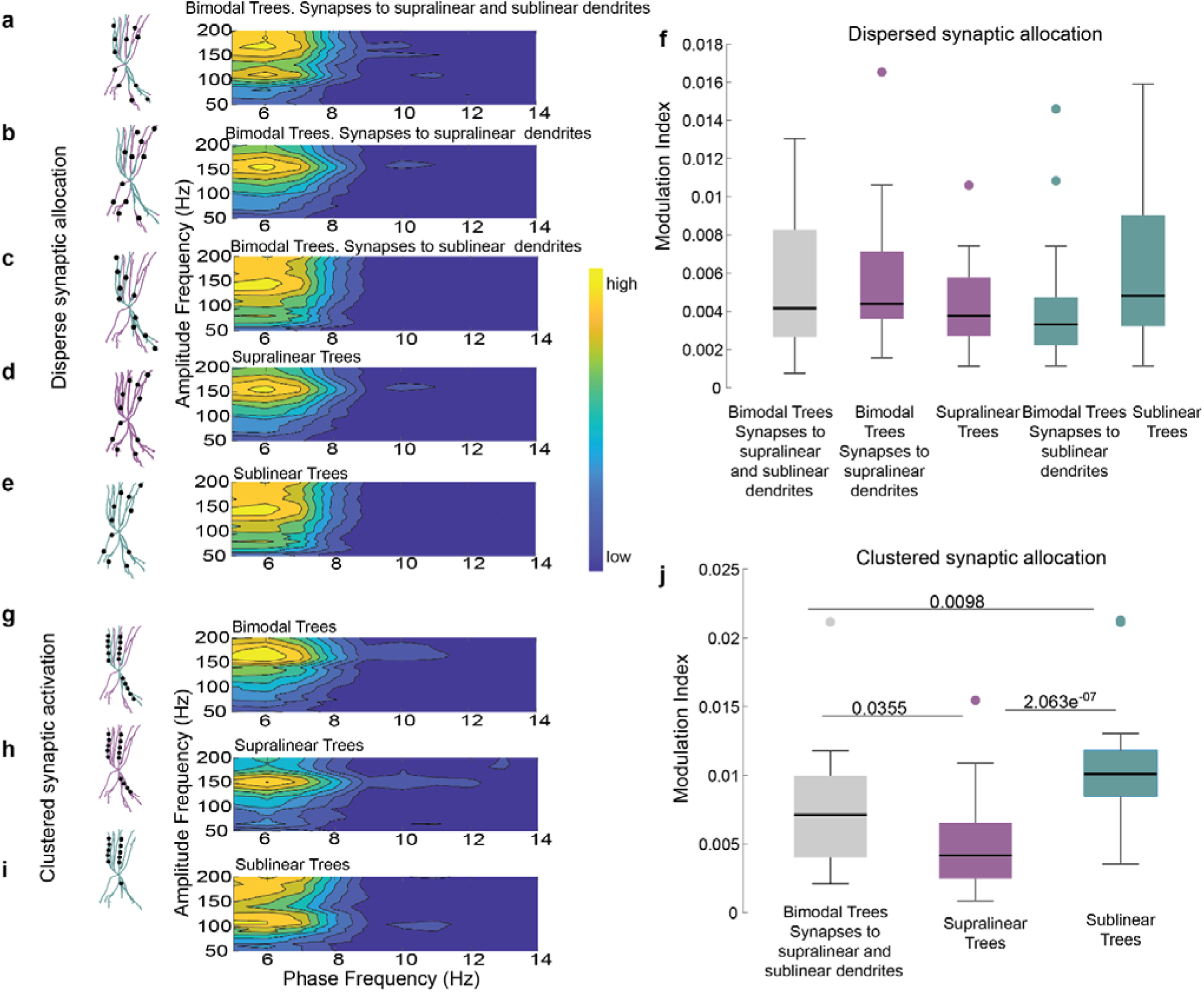
**Slow-Fast Oscillation Coupling in the Microcircuit Model Under Various Dendritic and Synaptic Configurations in FSBCs. a-e**. Representative comodulograms illustrating the slow-fast coupling for the protocols detailed in Figures 2 and S3. These visualizations provide insights into the phase-amplitude coupling dynamics under different synaptic and dendritic configurations. Data from 30 random simulation trials are represented. **f**. Coupling analysis shows that synaptic distribution in a dispersed configuration across bimodal, purely supralinear, or purely sublinear FSBC dendritic trees does not affect slow-fast oscillation coupling. **g-i**. Similar to a-e but showcasing comodulograms for a clustered synaptic arrangement (detailed in Figure S4), using the same number of synapses as in the dispersed experiments. **j.** MI indicates that slow-fast oscillation coupling decreases when synapses are clustered in purely sublinear dendritic trees compared to other configurations. Multigroup comparisons were performed using the Kruskal-Wallis test followed by a post-hoc correction for multiple comparisons for multi-group data with unequal variance. This statistical approach was chosen to accommodate the diversity in the data from 30 random simulation trials.

**Table S1.**
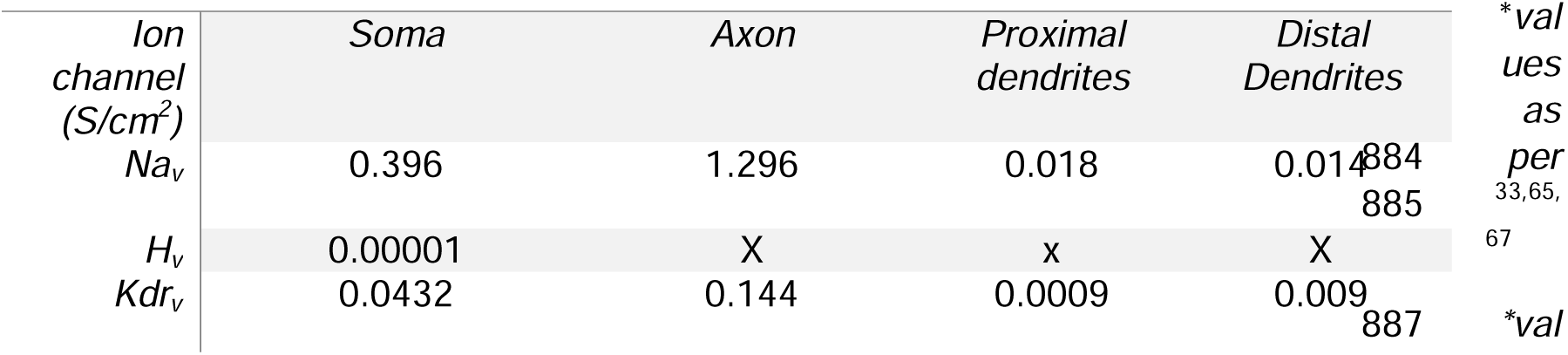
Active properties of Fast Spiking Basket Cell (FSBC) Models.

**Table S2.**
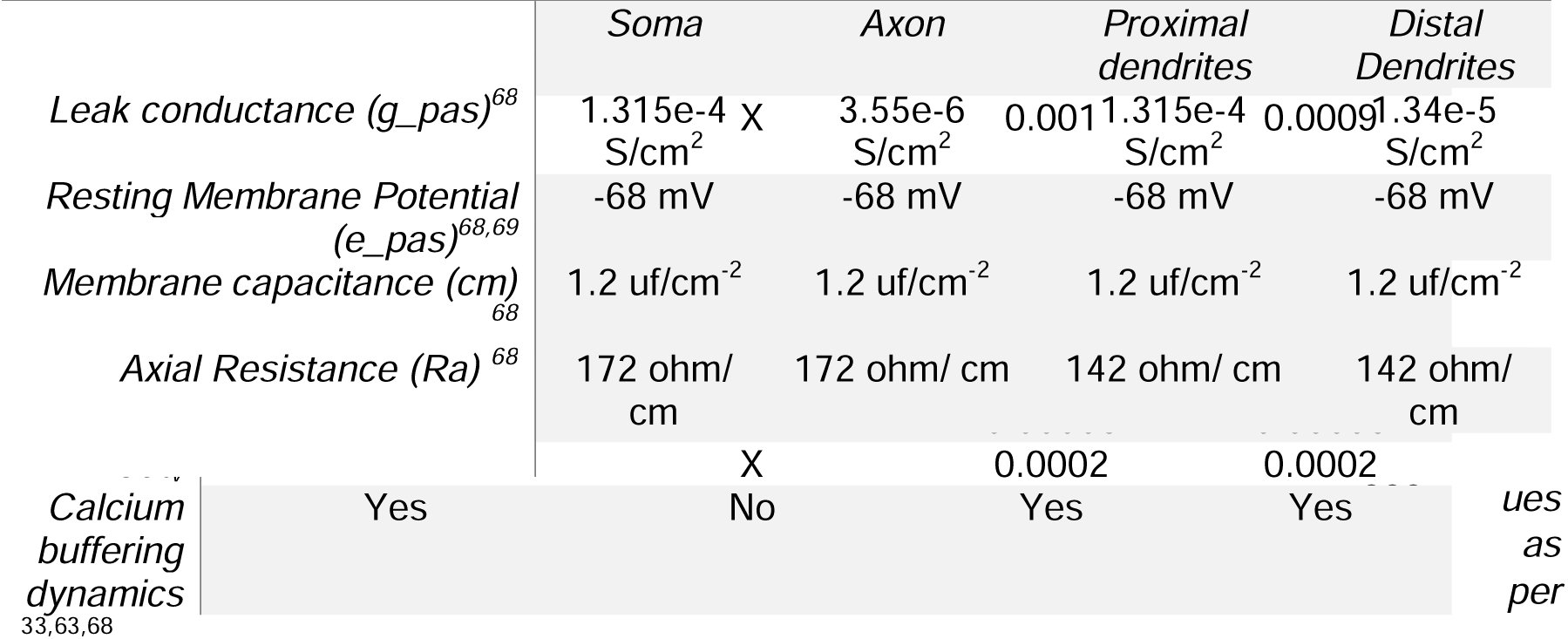
Passive properties of FSBCs.

**Table S3.**
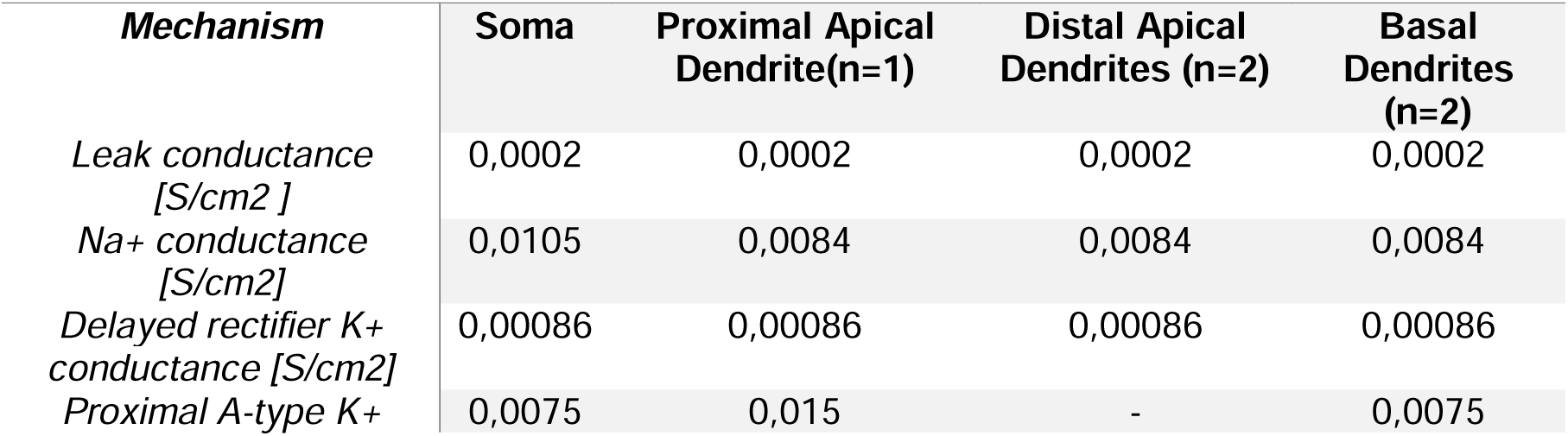
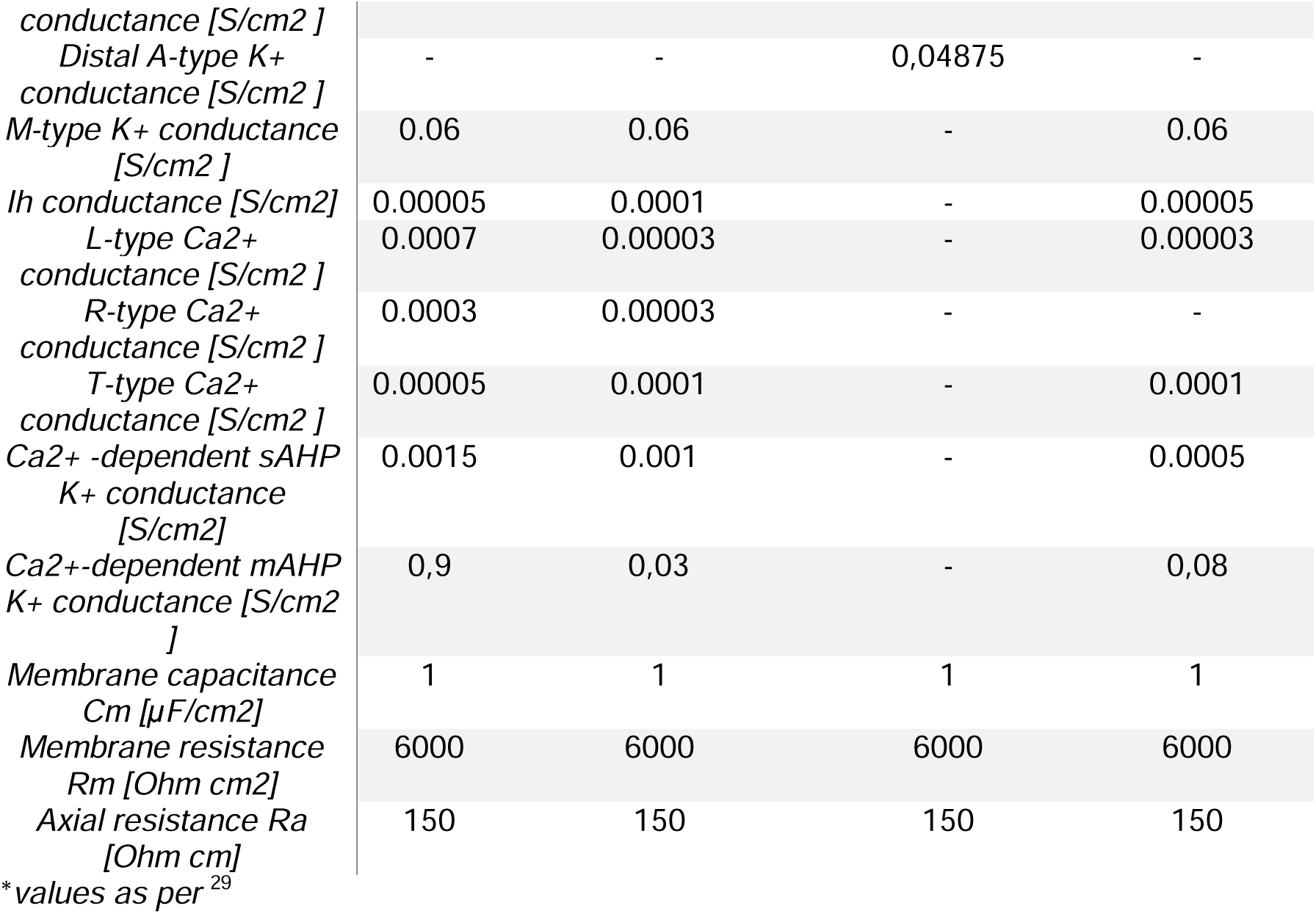
Passive Parameters and Active Conductance Values of the Pyramidal Cell (PC) Model.

**Table S4.**
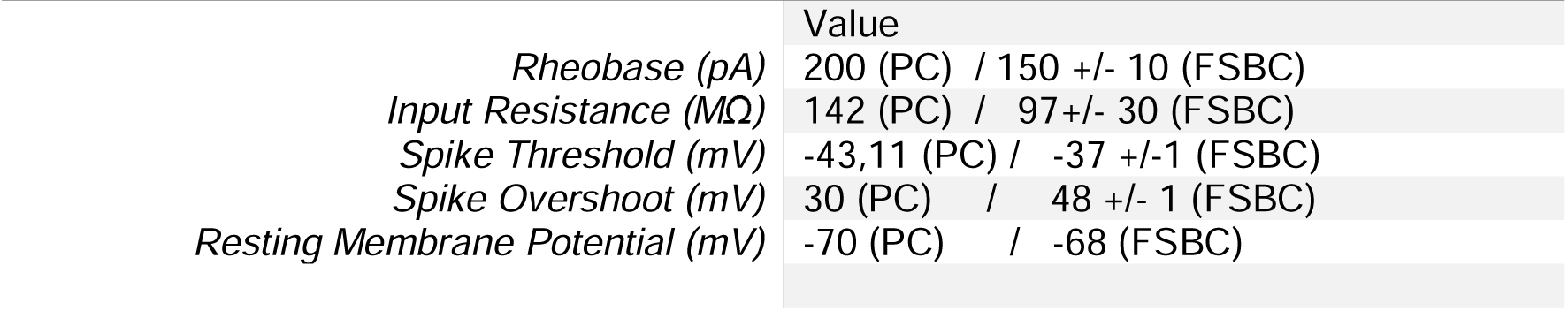
Electrophysiological properties of the PCs and FSBCs models.

**Table S5.**
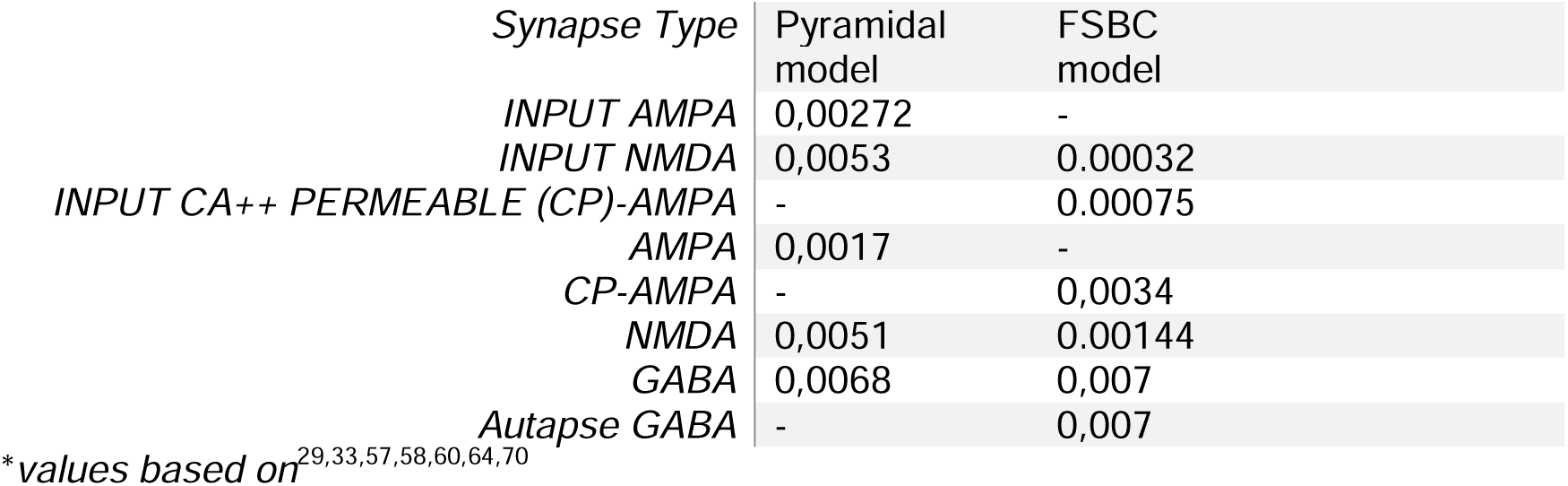
Synaptic conductance weight values of the PC and FSBC models.

## Glossary

Fast-Spiking Basket Cells (FSBCs): GABAergic inhibitory interneurons that primarily target the perisomatic region of pyramidal cells and other GABAergic neurons. Most FSBCs express parvalbumin (PV+) and are characterized by their ability to fire at high frequencies.
PV+ neuron: Parvalbumin-positive interneuron.
SST+ neuron: Somatostatin-positive interneuron.
VIP+ neuron: Vasoactive Intestinal Polypeptide-positive interneuron.
Bimodal Dendritic Trees: Dendritic trees in which different branches exhibit distinct nonlinear modes of excitatory synaptic integration: specifically, supralinear and sublinear. These integration modes coexist within the same neuron and are governed by differences in the morphological and/or active properties of individual dendritic branches.
EPSPs (Excitatory Postsynaptic Potential): Depolarizing electrical signals generated in the postsynaptic neuron upon the activation of excitatory synaptic input.
Dendritic Spikes: Local, nonlinear depolarizations in dendrites that reflect supralinear summation of excitatory postsynaptic potentials (EPSPs). Often modeled as a sigmoidal function of integrated EPSPs.
Memory Encoding: The cognitive process by which new information is initially acquired and stored—commonly referred to as learning.
Memory Consolidation: The process by which newly encoded information is stabilized and stored for long-term retrieval.
Memory Recall: The retrieval of previously encoded and consolidated information.
E/I Balance (Excitation/Inhibition Balance): The ratio of excitatory to inhibitory activity within a local circuit or network. An increase in inhibition lowers the E/I balance, while a reduction in inhibition raises it.
Neuronal Oscillations: Rhythmic fluctuations in electrical activity within the brain, typically measured via local field potentials (LFPs), that are associated with various behavioral and cognitive states.

