## Supplementary material for "Bimodal nonlinear dendrites in PV+ basket cells drive distinct memory-related oscillations": Figures S1-S6: Methods_SI_biorxiv_revised_Tzilivakietal.docx

Thata like probability formula (as per ^61^):

$$p(t)=(sin \left( 2.0*pi*\frac{f_{theta*spike}}{1000.0}+phi_{theta} \right)+1.0)/2.0$$

Where:

- spike = the spike time in msec (float)
- f_theta = theta-cycle frequency in Hz. (float)

For the simulations shown in **Figures 1-4** and **Supplementary Figures 4-6**, f_theta=4 Hz. For the sensitivity analysis simulations as shown in **Supplementary Figure 3**, we changed the input the theta frequency to f_theta=5Hz.

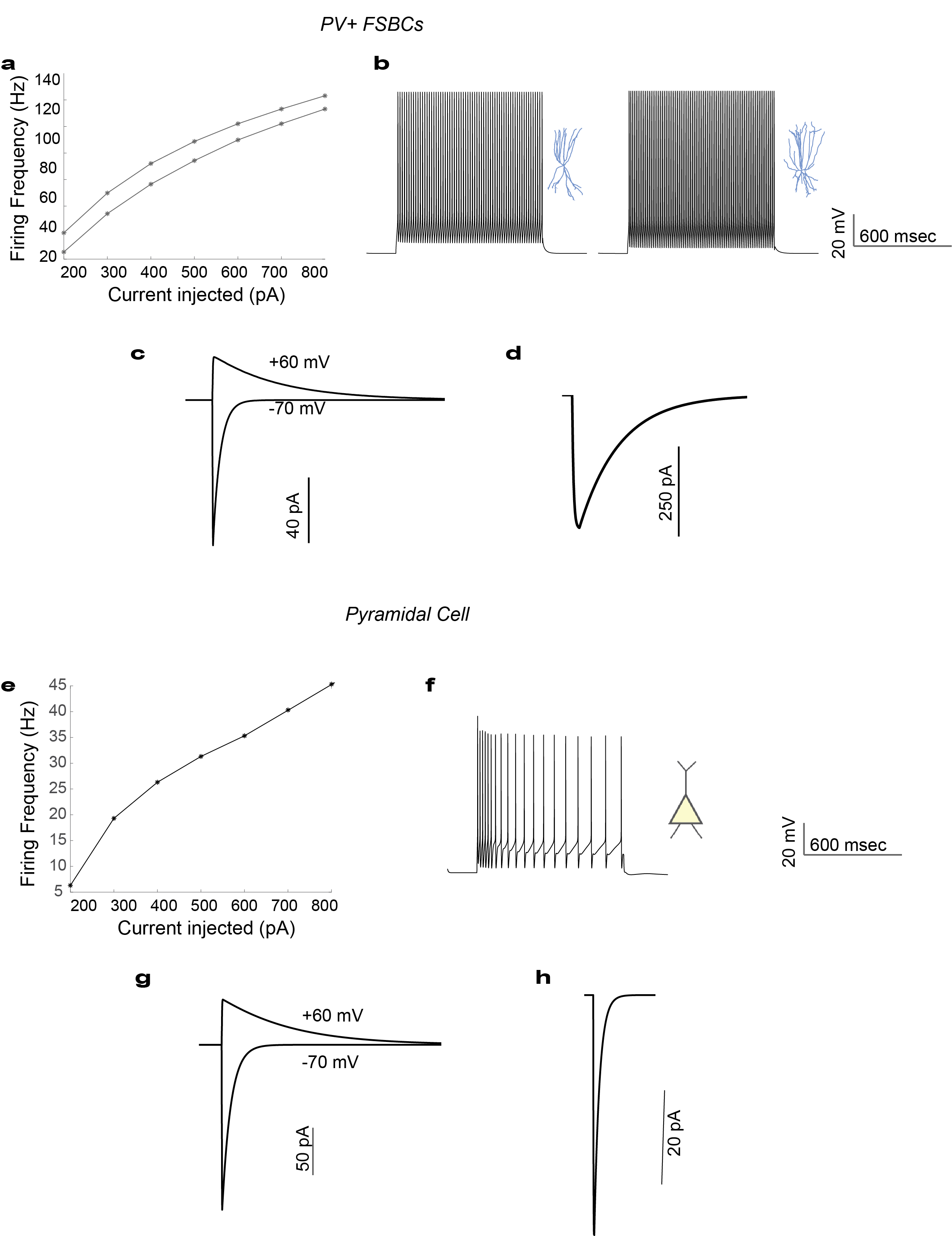
**Supplementary Information**

***Figure S1. Electrophysiological calibration and responses of the models. a-c.*** *FS BCs responses.* ***a.*** *Frequency- Injected current (FI) curve upon current clamp simulation in the cell bodies of the two multicompartmental FSBCs (1 sec duration).* ***b****. Firing profiles of the two FSBCs after depolarizing current injections in the cell bodies (300 pA, 1 sec). FSBCs exhibit the typical high-frequency firing pattern.* ***c.*** *CP-AMPA (−70 mV) and NMDA (+60 mV) currents upon stimulation as per ^60^. Traces represents the mean of the two multicompartmental FSBCs.* ***d.*** *A three-step voltage clamp of voltage changes from −70 mV to 10 mV (duration 1 msec) and back to −70 mV was used to produce inhibitory current. During the validation of this current, the reversal potential of Cl− was adjusted from −80 to −16 mV, in order to reproduce the experimental set up of ^63^, Mean trace of the two FSBCs responses.* ***e-h*** *Pyramidal Cell responses.* ***e.*** *FI curve of the Pyramidal Cell model same approach as in a.* ***f.*** *Firing profile of the Pyramidal Cell under current injection (300 pA for 1 sec) at the cell body. The model successfully represents the typical experimental phenotype shown in ^56^.* ***g.*** *AMPA (-70 mV voltage clamp) and NMDA (60 mV voltage clamp) currents of the Pyramidal Cell model mimic the experimental data of ^58^.* ***h.*** *Inhibitory current calibration based on experimental data from ^64^.*

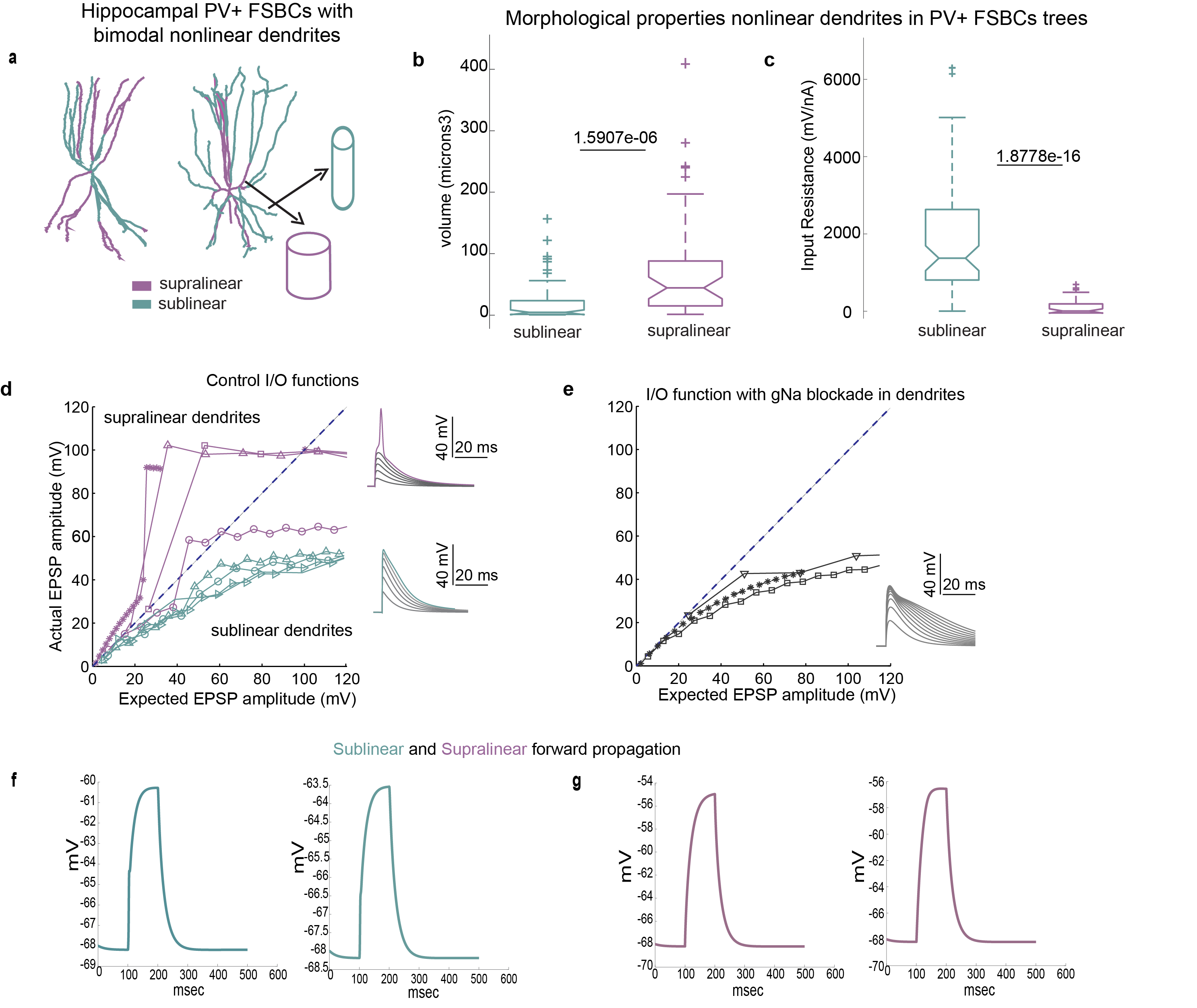

***Figure S2****.* ***Mechanisms of bimodal nonlinear dendritic integration in Multicompartmental Models of Hippocampal FSBCs. a****.Illustration of the morphological characteristics of supralinear and sublinear dendrites in bimodal FSBC models. Supralinear dendrites (purple) are larger, whereas sublinear dendrites (blue) are longer and thinner.* ***b-c****. Discriminative features between dendrite types: supralinear dendrites have larger volumes (b) and lower input resistance (c) compared to sublinear dendrites. Statistical significance was determined using the Mann–Whitney U test.* ***d-e****. Supralinear dendrites (purple) can generate local sodium-dependent spikes, whereas sublinear dendrites (blue) cannot (d). Blocking active sodium conductance in FSBC dendrites completely abolishes the supralinear mode (e).* ***f-g****. Forward propagation efficiency in bimodal nonlinear dendrites of FSBCs: Current injection (100 pA) at randomly selected dendrites and recording at the soma show that sublinear branches (f) propagate signals less effectively compared to supralinear branches (g). Panels a,d,e were adopted from ^33^*

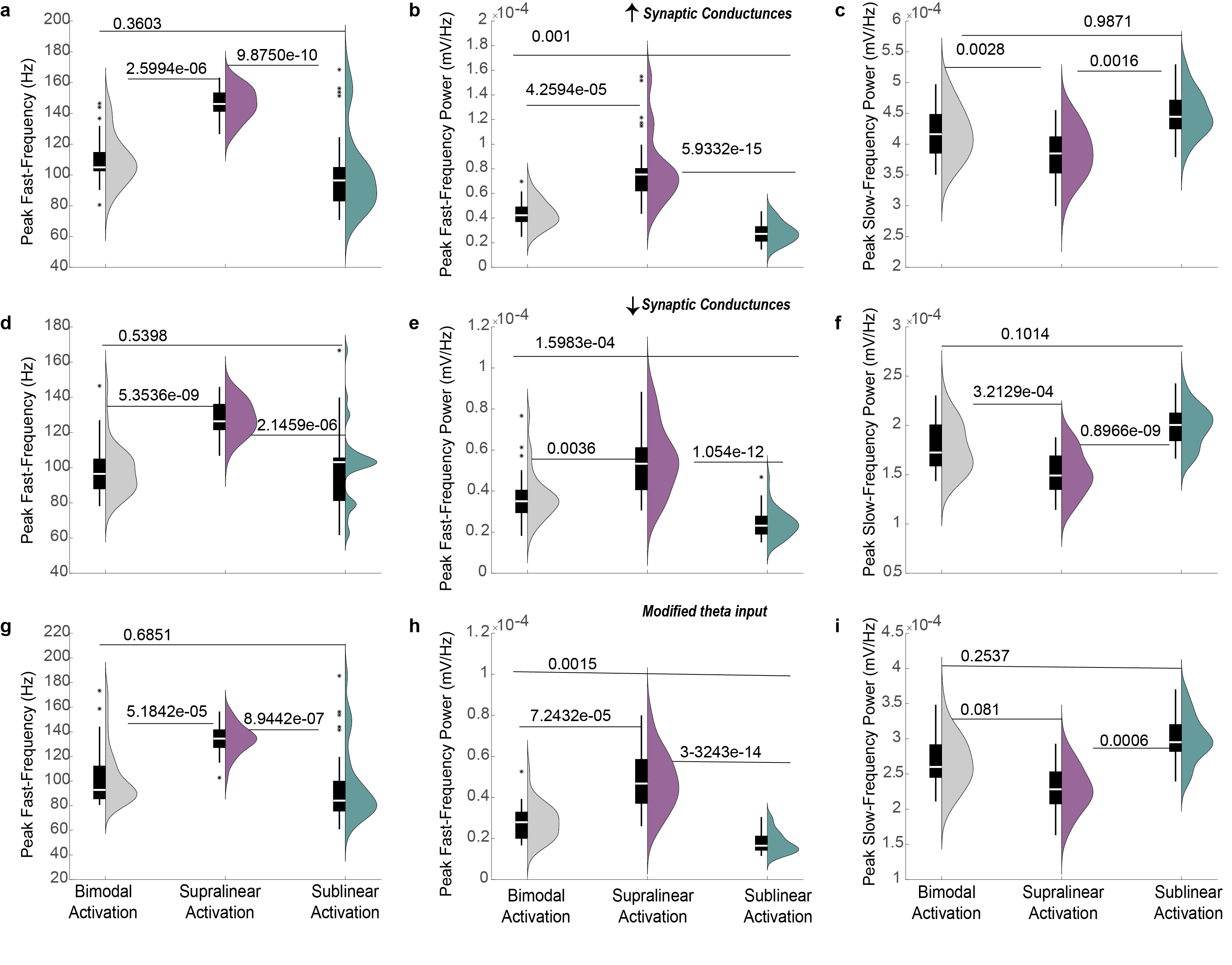

***Figure S3. Robustness/Sensitivity Analysis on Synaptic and Input Parameters. a-c****. 15% increase in the synaptic conductance values (applied to both input and network synapses for/from both PCs and FSBCs) do not alter the enhancement of fast peak frequency and power observed with supralinear activation.* ***d-f****. 15% reduction of the synaptic conductance values (for both input and network synapses involving PCs and FSBCs) also maintain the observed increase in fast peak frequency and power upon supralinear activation.* ***g-i****. Modifications to input parameters (input phase shifted 180°, peak frequency 5 Hz) similarly do not affect the enhancement of fast peak frequency and power driven by supralinear activation. Statistical analyses for multigroup comparisons were performed using the Kruskal-Wallis test, followed by a post-hoc correction for multiple comparisons.*

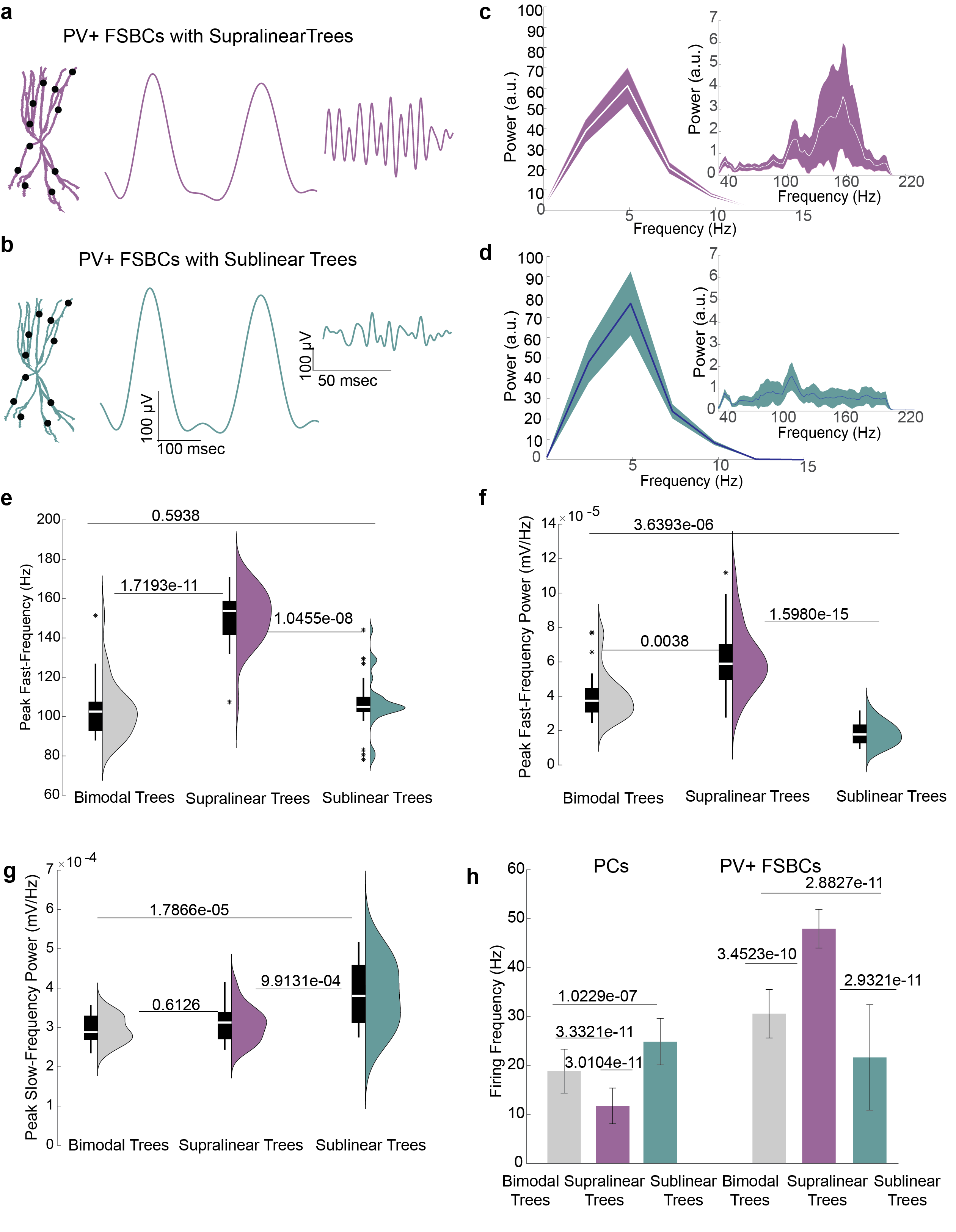

***Figure S4. Differential Modulation of Slow and Fast LFP Components by Supralinear and Sublinear FSBCs Dendritic Trees.*** ***a****. Activation of FSBCs equipped with purely supralinear dendritic trees, showcasing representative LFP traces bandpassed at slow (3-10 Hz) and high (30-200 Hz) frequencies.* ***b****. Similar to* ***a*** *but displaying activation of FSBCs with purely sublinear dendritic trees.* ***c-d****. Power Spectrum Density (PSD) plots of the LFP evoked when FSBCs are equipped with purely supralinear (c) or purely sublinear (d) dendritic trees, highlighting differences in frequency response.* ***e-g****. Comparative analysis of the peak fast-frequency (e) and peak power of fast (f) and slow (g) oscillations for FSBCs with bimodal (grey), supralinear (purple), or sublinear (blue) dendritic trees. Data are derived from 30 random simulation trials.* ***h****. Firing activity of the PCs and FSBCs populations within the microcircuit network across 30 random simulation trials. Activation of supralinear FSBC dendritic trees results in a decreased E/I balance compared to both bimodal (control) and sublinear trees.* *Statistical analyses for multigroup comparisons were conducted using the Kruskal-Wallis test followed by a post-hoc correction for multiple comparisons. Paired comparisons and p-values were calculated using the Mann-Whitney U test for data with unequal variance.*

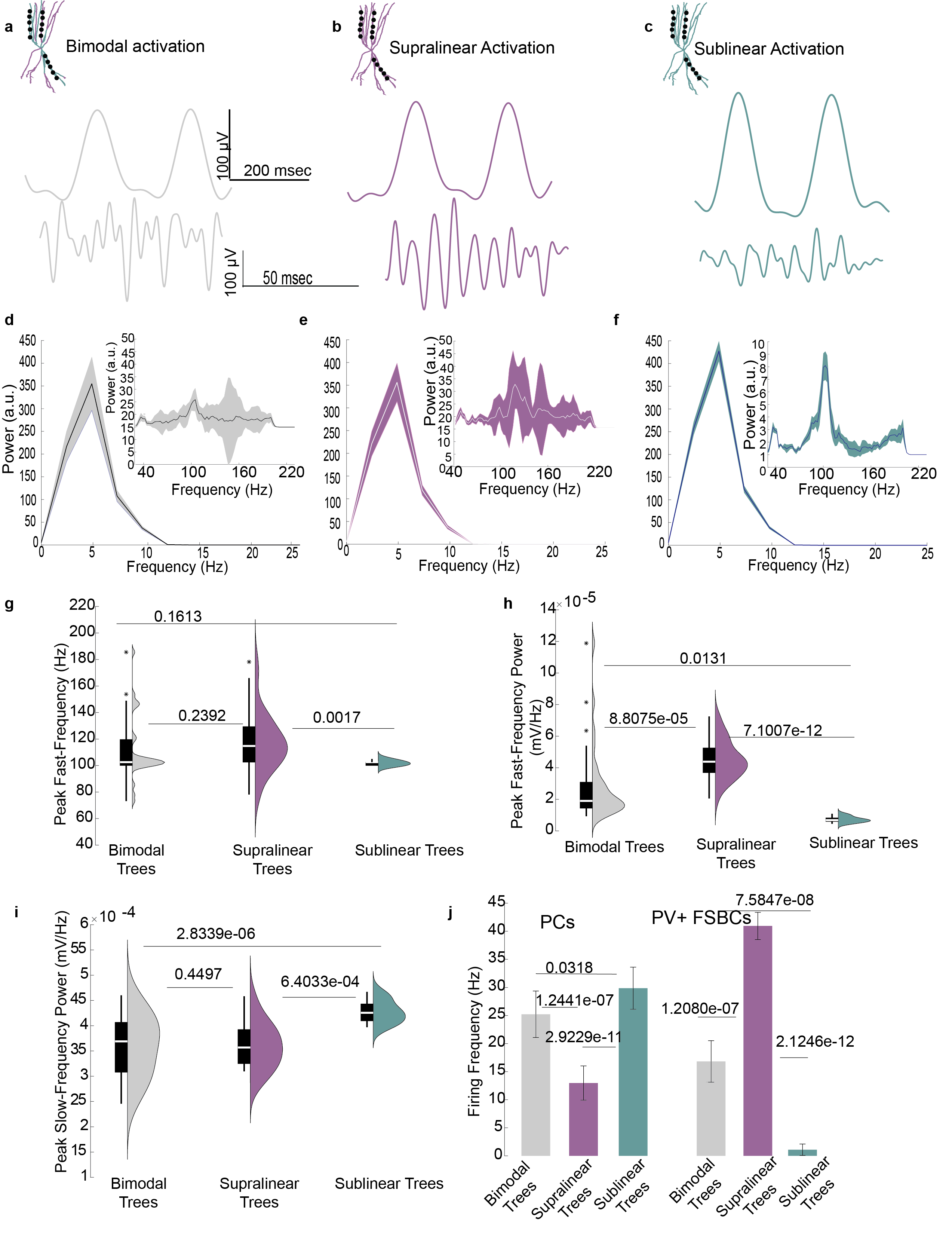

**Figure S5. Impact of Clustered Synaptic Activation on LFP and E/I Balance in Bimodal, Supralinear, and Sublinear FSBC Dendritic Configurations. a.** Activation of FSBCs with bimodal nonlinear dendrites (control configuration), showing representative LFP traces for slow (3-10 Hz) and fast (30-200 Hz) frequencies. Synapses are clustered in a few randomly chosen supralinear or sublinear branches. **b**. Activation of FSBCs with purely supralinear dendritic trees, showing LFP traces under clustered synaptic activation. **c**. Activation of FSBCs with purely sublinear dendritic trees, similar to conditions in a and b, illustrating the effect of clustered synaptic activation on LFP traces. **d-f**. Power Spectrum Density (PSD) plots for LFPs under clustered synaptic conditions in FSBCs with bimodal (d), supralinear (e), and sublinear (f) dendritic configurations. **g-i**. Comparison of peak frequency (g) and peak power for the fast LFP component (30-200 Hz) (h), and peak power for the slow LFP component (i), across dendritic configurations. **j**. Firing activity of PC and FSBC populations in the microcircuit network from 30 random simulation trials. Clustering in supralinear FSBC dendritic trees decreases the E/I balance in the network compared to bimodal (control) or sublinear trees. Statistical analyses for multigroup comparisons were conducted using the Kruskal-Wallis test followed by a post-hoc correction for multiple comparisons. Paired comparisons and p-values were calculated using the Mann-Whitney U test for data with unequal variance.

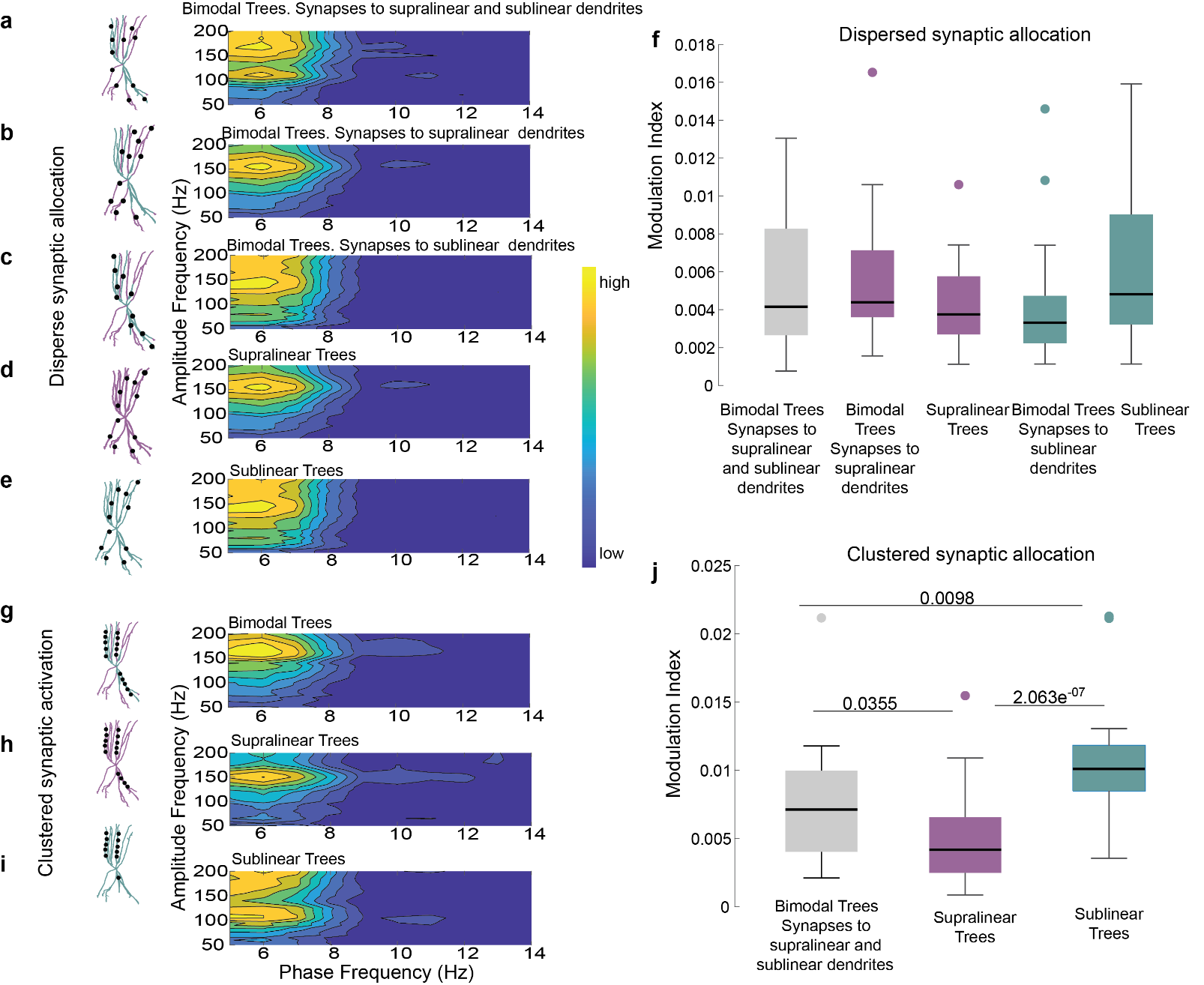

***Figure S6. Slow-Fast Oscillation Coupling in the Microcircuit Model Under Various Dendritic and Synaptic Configurations in FSBCs. a-e****. Representative comodulograms illustrating the slow-fast coupling for the protocols detailed in Figures 2 and S3. These visualizations provide insights into the phase-amplitude coupling dynamics under different synaptic and dendritic configurations. Data from 30 random simulation trials are represented.* ***f****. Coupling analysis shows that synaptic distribution in a dispersed configuration across bimodal, purely supralinear, or purely sublinear FSBC dendritic trees does not affect slow-fast oscillation coupling.* ***g-i****. Similar to a-e but showcasing comodulograms for a clustered synaptic arrangement (detailed in Figure S4), using the same number of synapses as in the dispersed experiments.* ***j.*** *MI indicates that slow-fast oscillation coupling decreases when synapses are clustered in purely sublinear dendritic trees compared to other configurations. Multigroup comparisons were performed using the Kruskal-Wallis test followed by a post-hoc correction for multiple comparisons for multi-group data with unequal variance. This statistical approach was chosen to accommodate the diversity in the data from 30 random simulation trials.*

| **Table S1. Active properties of Fast Spiking Basket Cell (FSBC) Models** | | | | |
| --- | --- | --- | --- | --- |
| Ion channel (S/cm^2^) | *Soma* | *Axon* | *Proximal dendrites* | *Distal Dendrites* |
| Na_v_ | 0.396 | 1.296 | 0.018 | 0.014 |
| H_v_ | 0.00001 | X | x | X |
| Kdr_v_ | 0.0432 | 0.144 | 0.0009 | 0.009 |
| Kslow_v_ | 0.000725 | X | x | X |
| Kct_v_ | 0.0001 | X | x | X |
| Kca_v_ | 0.02 | X | x | X |
| Ka_v_ (proximal type) ^65^ | 0.0032 | X | 0.001 | 0.0009 |
| Ka_v_ (distal type) ^66^ | x | X | x | 0.00216 |
| Cal_v_ | x | X | 0.00003 | 0.00003 |
| Can_v_ | x | X | 0.00003 | 0.00003 |
| Cat_v_ | x | X | 0.0002 | 0.0002 |
| Calcium buffering dynamics | Yes | No | Yes | Yes |

**values as per* ^33,65,67^

| **Table S2. Passive properties of FSBCs** | | | | |
| --- | --- | --- | --- | --- |
|  | *Soma* | *Axon* | *Proximal dendrites* | *Distal Dendrites* |
| Leak conductance (g_pas)^68^ | 1.315e-4 S/cm^2^ | 3.55e-6 S/cm^2^ | 1.315e-4 S/cm^2^ | 1.34e-5 S/cm^2^ |
| Resting Membrane Potential (e_pas)^68,69^ | -68 mV | -68 mV | -68 mV | -68 mV |
| Membrane capacitance (cm) ^68^ | 1.2 uf/cm^-2^ | 1.2 uf/cm^-2^ | 1.2 uf/cm^-2^ | 1.2 uf/cm^-2^ |
| Axial Resistance (Ra) ^68^ | 172 ohm/ cm | 172 ohm/ cm | 142 ohm/ cm | 142 ohm/ cm |

**values as per* ^33,63,68^

| **Table S3. Passive Parameters and Active Conductance Values of the Pyramidal Cell (PC) Model** | | | | |
| --- | --- | --- | --- | --- |
| **Mechanism** | **Soma** | **Proximal Apical Dendrite(n=1)** | **Distal Apical Dendrites (n=2)** | **Basal Dendrites (n=2)** |
| Leak conductance [S/cm2 ] | 0,0002 | 0,0002 | 0,0002 | 0,0002 |
| Na+ conductance [S/cm2] | 0,0105 | 0,0084 | 0,0084 | 0,0084 |
| Delayed rectifier K+ conductance [S/cm2] | 0,00086 | 0,00086 | 0,00086 | 0,00086 |
| Proximal A-type K+ conductance [S/cm2 ] | 0,0075 | 0,015 | - | 0,0075 |
| Distal A-type K+ conductance [S/cm2 ] | - | - | 0,04875 | - |
| M-type K+ conductance [S/cm2 ] | 0.06 | 0.06 | - | 0.06 |
| Ih conductance [S/cm2] | 0.00005 | 0.0001 | - | 0.00005 |
| L-type Ca2+ conductance [S/cm2 ] | 0.0007 | 0.00003 | - | 0.00003 |
| R-type Ca2+ conductance [S/cm2 ] | 0.0003 | 0.00003 | - | - |
| T-type Ca2+ conductance [S/cm2 ] | 0.00005 | 0.0001 | - | 0.0001 |
| Ca2+ -dependent sAHP K+ conductance [S/cm2] | 0.0015 | 0.001 | - | 0.0005 |
| Ca2+-dependent mAHP K+ conductance [S/cm2 ] | 0,9 | 0,03 | - | 0,08 |
| Membrane capacitance Cm [μF/cm2] | 1 | 1 | 1 | 1 |
| Membrane resistance Rm [Ohm cm2] | 6000 | 6000 | 6000 | 6000 |
| Axial resistance Ra [Ohm cm] | 150 | 150 | 150 | 150 |

**values as per* ^29^

| **Table S4. Electrophysiological properties of the PCs and FSBCs models** | |
| --- | --- |
|  | Value |
| Rheobase (pA) | 200 (PC) / 150 +/- 10 (FSBC) |
| Input Resistance (MΩ) | 142 (PC) / 97+/- 30 (FSBC) |
| Spike Threshold (mV) | -43,11 (PC) / -37 +/-1 (FSBC) |
| Spike Overshoot (mV) | 30 (PC) / 48 +/- 1 (FSBC) |
| Resting Membrane Potential (mV) | -70 (PC) / -68 (FSBC) |

| **Table S5. Synaptic conductance weight values of the PC and FSBC models.** | | |
| --- | --- | --- |
| Synapse Type | Pyramidal model | FSBC  model |
| INPUT AMPA | 0,00272 | - |
| INPUT NMDA | 0,0053 | 0.00032 |
| INPUT CA++ PERMEABLE (CP)-AMPA | - | 0.00075 |
| AMPA | 0,0017 | - |
| CP-AMPA | - | 0,0034 |
| NMDA | 0,0051 | 0.00144 |
| GABA | 0,0068 | 0,007 |
| Autapse GABA | - | 0,007 |
